# Mechanistic insights into dynamic mutual regulation of USP14 and proteasome

**DOI:** 10.1101/2021.09.15.460436

**Authors:** Shuwen Zhang, Shitao Zou, Deyao Yin, Daniel Finley, Zhaolong Wu, Youdong Mao

**Affiliations:** State Key Laboratory for Mesoscopic Physics, School of Physics, Peking University, Beijing, China; Center for Quantitative Biology, Peking University, Beijing, China; Department of Cell Biology, Harvard Medical School, Boston, MA, USA; National Biomedical Imaging Center, Peking University, Beijing, China

## Abstract

Proteasomal degradation of ubiquitylated proteins is sophisticatedly regulated at multiple levels^1–3^. A primary regulatory checkpoint is the removal of ubiquitin chains from substrates by the deubiquitylating enzyme USP14 that associates reversibly with the proteasome. How USP14 is activated and regulates the proteasome function remains unknown^4–7^. Here we report cryo-electron microscopy (cryo-EM) structures of human USP14 in complex with the 26S proteasome in nine conformational states at 3.0-3.6 Å resolution, captured during polyubiquitylated protein degradation. Time-resolved cryo-EM analysis of the conformational continuum revealed two parallel pathways of proteasome state transitions induced by USP14 and captured transient conversion of substrate-engaged intermediates into substrate-inhibited intermediates. On the substrate-engaged pathway, USP14 activation allosterically reprograms conformational landscape of the AAA-ATPase motor and stimulates opening of the core particle gate^8–10^, enabling observation of a near-complete cycle of asymmetric ATP hydrolysis around the ATPase ring during processive substrate unfolding. Dynamic USP14-ATPase interactions decouple the ATPase activity from RPN11-catalysed deubiquitylation^11–13^ and kinetically introduce three regulatory checkpoints on the proteasome, at the steps of ubiquitin recognition, substrate translocation initiation and ubiquitin chain recycling. These findings provide unprecedented insights into the complete functional cycles of USP14-regulated proteasome and of USP14 activation-deubiquitylation-disassembly and establish mechanistic foundations for USP14-targeted therapeutic discovery.

---

Majority of cellular proteins are targeted to the proteasome for degradation by ubiquitylation pathways, which regulates nearly all major aspects of cellular processes in eukaryotes^1, 10^. The proteasome holoenzyme is assembled by a cylindrical 20S core particle (CP) capped with one or two 19S regulatory particles (RPs), each consisting of the lid and base subcomplexes^1, 10^. The ring-like heterohexameric motor of ATPases-associated-with-diverse-cellular-activities (AAA) ATPase in the base subcomplex regulates substrate engagement, deubiquitylation and translocation in the proteasome via multiple modes of coordinated ATP hydrolysis^10^. The proteasome is also sophisticatedly regulated by numerous proteins that reversibly associate with it^3^. Little is known about the mechanisms of such extrinsic regulations of the proteasome.

Human ubiquitin-specific protease 14 (USP14) is one of the three proteasome-associated deubiquitinating enzymes (DUBs)^2^ and a potential therapeutic target for major neurodegenerative diseases^6^. In contrast to the stoichiometric DUB subunit RPN11, USP14 is the most frequently observed DUB that reversibly associates with the human proteasome^4–7^. USP14 retains a low level of basal DUB activity in isolation and is prominently activated upon its incorporation into the proteasome^4–7^. On the proteasome, USP14 appears to remove each supernumerary ubiquitin chain on a substrate en bloc until a single chain remains^7^. It was hypothesized that USP14 inhibits proteasomal degradation by both catalytically removing ubiquitin chains prematurely and non-catalytically delaying proteasomal degradation^14, 15^. Neither mechanism is structurally defined and well understood. The kinetic competition between USP14 and the proteasome provides a critical layer of regulation of proteasome activity. Unlike the DUB activity of RPN11 that is allosterically coupled to the proteasomal ATPase activities, USP14 appears to trim ubiquitin chains independently of ATP-driven substrate engagement. Curiously, the proteasomal ATPase activity seems to be stimulated by USP14 in the presence of ubiquitylated substrates^16–20^. The molecular mechanism underlying the proteasome-mediated USP14 activation and its reciprocal regulation of the proteasome function remains elusive.

Previous cryogenic electron microscopy (cryo-EM) studies at low resolution have revealed the approximate location of USP14 or its yeast orthologue Ubp6 in the proteasome^18,21,22^. However, the insufficient resolution of USP14/Ubp6-assigned cryo-EM densities, the unclear ubiquitin-like (UBL) domain of USP14 and the absence of polyubiquitylated substrate interactions in these structural studies preclude understanding of USP14-mediated proteasome regulation at atomic level. Particularly, the full-length USP14 structure has not been determined to date. Here we describe high-resolution cryo-EM structures of human full-length USP14 in complex with functional proteasome in the act of substrate degradation. Time-resolved cryo-EM analysis of USP14-loaded proteasomal degradation portrays a dynamic picture of USP14-mediated proteasome regulation and visualizes a series of human proteasome intermediates that are uniquely induced by USP14. Structure-guided mutagenesis experiments substantiate the functional importance of key structural elements at dynamic USP14-proteasome interfaces. Our comprehensive analysis provides unprecedented insights into the mechanism of allosteric ‘tug-of-war’ between USP14 and the proteasome for deciding substrate fate.

## Visualizing intermediates of USP14-regulated proteasome

To prepare a substrate-engaged USP14-proteasome complex, we separately purified human USP14, RPN13 and the USP14-free 26S proteasome. The purified USP14 and RPN13 in excessive stoichiometry were mixed with the USP14-free proteasome to maximize the USP14 presence in the proteasome (Fig. 1, Extended Data Fig. 1). We used polyubiquitylated Sic1^PY^ (Ub_*n*_-Sic1^PY^) as a model substrate in this study, which has been previously used to characterize USP14 function in the proteasome^7^. To optimize sample preparation condition for maximally capturing intermediate states of the USP14-bound proteasome during substrate processing, we first collected cryo-EM data on the samples prepared in a time-dependent manner, by cryo-plunging in 1, 5 and 10 minutes after mixing Ub_*n*_-Sic1^PY^ with the USP14-proteasome in the presence of 1 mM ATP at 10 °C. 3D classification of cryo-EM data indicated that intermediate states are approximately maximized in 1 min after substrate addition (Fig. 1e). Thus, we further collected two large datasets with and without using ATP-to-ATPγS exchange to quench the ATPase activity in 1 minute after mixing Ub_*n*_-Sic1^PY^ with the USP14-proteasome at 10 °C (see Methods)^10^.

**Figure 1.**
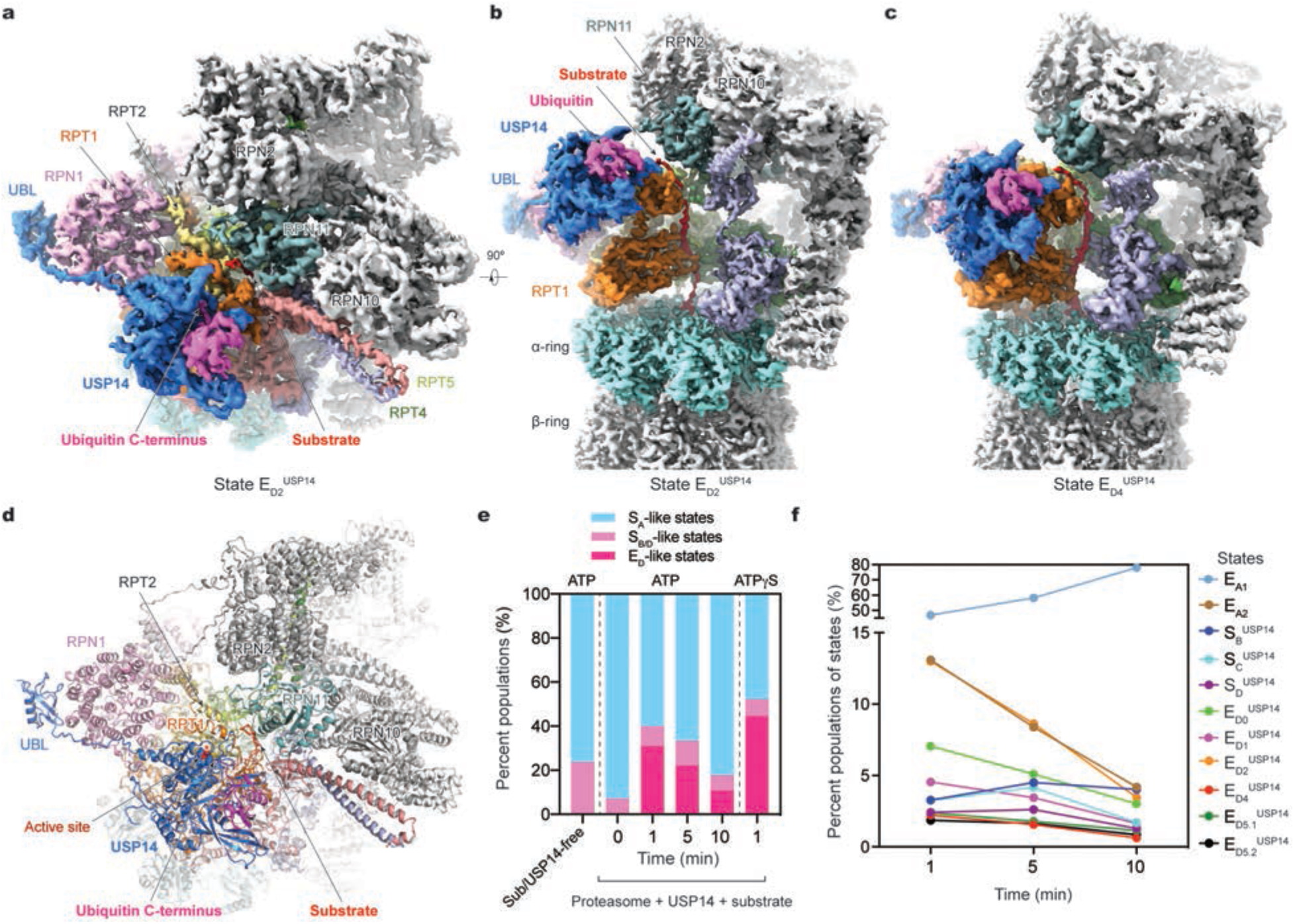
Cryo-EM structures of the substrate-engaged USP14-proteasome complexes in representative states and time-resolved cryo-EM analysis of the conformational continuum of USP14-proteasome. **a**, High-resolution cryo-EM density map of the substrate-engaged USP14-proteasome complex in state E_D2_^USP14^, viewed from the top perspective approximately along the axis of the AAA-ATPase ring, where both the USP14-bound ubiquitin and ATPase-engaged substrate are visible. **b**, Side-view of the cryo-EM map of the substrate-engaged USP14-proteasome complex in state E_D2_^USP14^ that is rotated from the view shown in panel (**a**). **c**, Side-view of the high-resolution cryo-EM map of the substrate-engaged USP14-proteasome complex in state E_D4_^USP14^. Compared to panel (**b**), USP14 is rotated about 30° to dock onto the AAA domain of RPT1. For clarify of the substrate density inside the AAA-ATPase motor, the density of RPT5 is omitted in both (**b**) and (**c**). **d**, The atomic model of state E_D2_^USP14^ viewed from the same perspective as the cryo-EM density in panel (**a**). **e**, Kinetic changes of overall particle populations of S_B/D_-like and E_D_-like states versus S_A_-like states obtained by using time-resolved cryo-EM sample preparation and 3D classification. S_A_-like states include E_A1_ and E_A2_. S_B/D_-like states include S_B_^USP14^, S_C_^USP14^ and S_D_^USP14^. E_D_-like states include E_D4_^USP14^, E_D5.1_^USP14^, E_D5.2_^USP14^, E_D0_^USP14^, E_D1_^USP14^ and E_D2_^USP14^. The control was measured on the previously reported data of substrate-free, USP14-free proteasome^8^. **f**, Kinetic changes of the particle populations of 11 coexisting conformational states observed on the same cryo-EM samples made at three different time points after mixing the substrate with the USP14-proteasome complex in the presence of 1 mM ATP at 10 °C. Notably, the three substrate-inhibited intermediate states (S_B_^USP14^, S_C_^USP14^ and S_D_^USP14^) reach their maximal populations at around 5 min, in contrast to the six substrate-engaged states (E_D4_^USP14^, E_D5.1_^USP14^, E_D5.2_^USP14^, E_D0_^USP14^, E_D1_^USP14^ and E_D2_^USP14^) that reach their maximal populations at approximately 1 min.

Focused 3D classification allowed us to determine six substrate-engaged conformational states (designated E_D0_^USP14^, E_D1_^USP14^, E_D2_^USP14^, E_D4_^USP14^, E_D5.1_^USP14^ and E_D5.2_^USP14^) and three substrate-free states (designated S_B_^USP14^, S_C_^USP14^ and S_D_^USP14^) of the USP14-bound proteasome at nominal resolutions of 3.0-3.6 Å (Fig. 1a–c, Extended Data Figs. 2, 3, Extended Data Table 1). The cryo-EM density of USP14 in state E_D2_^USP14^ is of sufficient quality to allow for atomic modeling of a full-length USP14 structure (Extended Data Fig. 4c, d). In addition, we also observed two coexisting states E_A1_ and E_A2_ that show no visible USP14 density except for the UBL-resembling density on RPN1 and that are virtually identical to their previously reported structures^10^ (Extended Data Fig. 4). Our cryo-EM analysis suggests that ATP-to-ATPγS exchange enriches states S_C_^USP14^, E_D2_^USP14^ and E_D4_^USP14^ but reduces states S_B_^USP14^, S_D_^USP14^ and E_D0_^USP14^ with an overall agreement with the ATP-only condition on the characterization of the conformational continuum (Extended Data Fig. 1j).

The six substrate-engaged states of USP14-proteasome captured sequential intermediate conformations during processive substrate unfolding and translocation, which are compatible with the hand-over-hand translocation model^10, 23^ (Extended Data Fig. 5, Extended Data Table 2). All these states exhibit an open CP gate, with five RPT C-tails inserted into the α-pockets of the CP (Extended Data Fig. 6a), indicating that these are translocation-competent states. While the overall proteasome conformations of states E_D0_^USP14^, E_D1_^USP14^ and E_D2_^USP14^ closely resemble those of USP14-free, substrate-engaged states E_D0.2_, E_D1.2_ and E_D2_, only the AAA-ATPase components in states E_D4_^USP14^, E_D5.1_^USP14^ and E_D5.2_^USP14^ are comparable to those of USP14-free states EB, EC1 and EC2 with defined differences, respectively^10, 25^ (Extended Data Figs. 5, 6b). For instance, the pore-1 loop of RPT6 in state E_D4_^USP14^ has considerably moved up toward the substrate as compared to that in state E_B_ (Extended Data Fig. 5c).

Like the USP14-free, substrate-free state S_B_, RPN11 blocks the substrate entrance at the oligonucleotide- or oligosaccharide-binding (OB) ring of the AAA-ATPase motor in states S_B_^USP14^, S_C_^USP14^ and S_D_^USP14^, indicating that these states sterically inhibit substrate insertion into the AAA-ATPase channel. The AAA-ATPase motors of these substrate-inhibited states all adopt an S_A_-like conformation^8, 9^. However, the ATPase-CP interfaces of state S_B_^USP14^, S_C_^USP14^ and S_D_^USP14^ resemble those of state E_A_, E_C_ and E_D_, where two, four and five RPT C-tails are inserted into the α-pockets, respectively^10^. Thus, the CP gate remains closed in S_B_^USP14^ and S_C_^USP14^, but is open in S_D_^USP14^ (Extended Data Fig. 6a). Taken together, USP14 binding appears to induce a greater degree of proteasome dynamics and expand its conformational space to sample both substrate-engaged and substrate-inhibited states.

## Time-dependent changes of conformational continuum

Figure 1f plots the time-dependent changes of the populations of all coexistent conformational states (Fig. 1f). Notably, the substrate-engaged and substrate-inhibited intermediates reach their maximal populations around 1 and 5 minutes, respectively, after mixing Ub_*n*_-Sic1^PY^ with the USP14-proteasome. At 10 minutes after substrate addition to the USP14-proteasome, the population of S_B_^USP14^ became the largest among the USP14-bound states. The population difference between S_B_^USP14^ and other inhibitory states (S_C_^USP14^ and S_D_^USP14^) was also widened over time. These kinetic features indicate that the substrate-engaged intermediates were converted into the substrate-inhibited states upon termination of substrate translocation, and that S_C_^USP14^ and S_D_^USP14^ changed into S_B_^USP14^ in the late stage of substrate processing. Indeed, structural comparison between S_D_^USP14^ and E_D4_^USP14^, which both have an open CP gate, indicates a highly comparable AAA-ATPase motor conformation, suggesting that S_D_^USP14^ is perhaps an intermediate state connecting E_D4_^USP14^ with other inhibitory states (Extended Data Fig. 6b).

Despite in-depth 3D classification, we did not observe the proteasome states E_B_ and E_C_ in all experimental conditions that represent RPN11-mediated deubiquitylation and translocation initiation prior to the CP gate opening, respectively^10^. This indicates that USP14 prevents the proteasome from assuming the conformation of RPN11-catalyzed deubiquitylation. Notably, the coexisting USP14-invisible state E_A2_, in which RPN11 is bound with substrate-conjugated ubiquitin, was observed at all time points to be on the population level similar to that of state E_D2_^USP14^, the most popular among the substrate-engaged intermediates (Fig. 1f). In contrast to the RPN11-bound ubiquitin in state E_A2_, one ubiquitin moiety binds the C-terminal catalytic USP domain with no ubiquitin on RPN11 in all USP14-loaded states (Fig. 1a–d, Extended Data Fig. 4), suggesting that USP14 and RPN11 do not bind ubiquitin simultaneously.

## Dynamic USP14-proteasome interactions

The human USP14 is composed of 494 amino acids in full length and features a 9-kDa UBL domain at its N-terminus, followed by a 43-kDa USP domain via a flexible linker region of 23 amino acids. The UBL domain interfaces RPN1 via a hydrophobic patch centered on residue Leu70 structurally homologous to the Ile44 surface of ubiquitin (Fig. 2a). It contacts the RPN1 T2 site composed of residues Asp423, Leu426, Asp430, Tyr434, Glu458, Asp460 and Leu465 on two adjacent helix-loop regions, agreeing with previous findings^24, 25^. The N-terminal stretch of the linker (residues Ala77 to Phe88) in USP14 appears to bind the ridge of the toroid domain of RPN1 (Fig. 2a).

**Figure 2.**
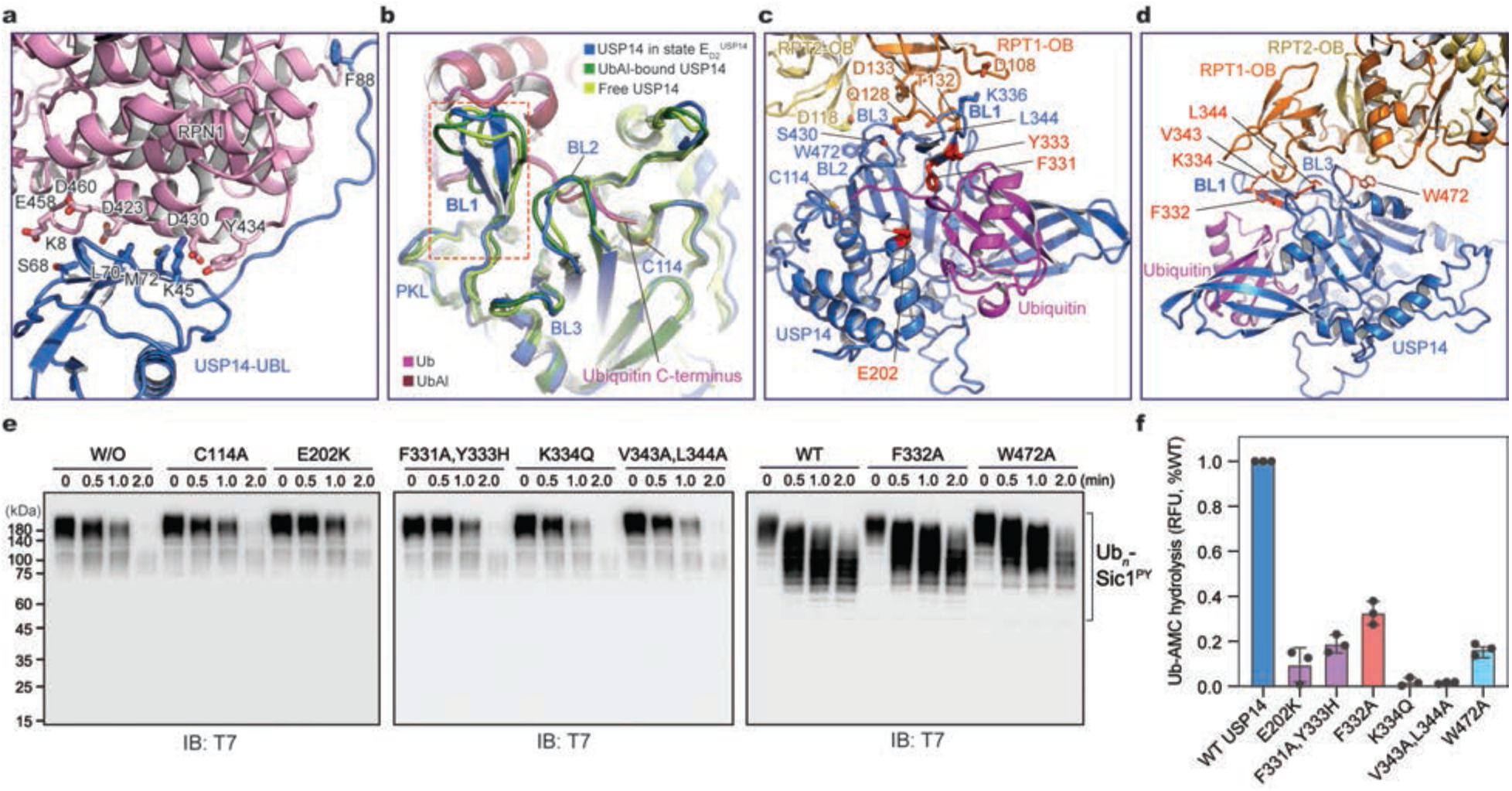
Structural basis of proteasome-mediated activation of USP14. **a**, Closeup view of the interaction between the UBL domain of USP14 and the T2 site of RPN1 in the proteasome of state E_D2_^USP14^. The acidic residues on the RPN1 T2 site interacting with the USP14 UBL domain are shown as stick representation and labeled. Several hydrophobic and basic residues on the UBL domain are shown as stick representation, among which Phe88 appears to anchor N-terminal part of the linker region on the RPN1 toroid domain. **b**, Structural comparison of the blocking loops by superimposing the USP14 structure in the proteasome of state E_D2_^USP14^ with two crystal structures of USP14 in its isolated form (PDB ID 2AYN) and in complex with UbAl (PDB ID 2AYO)^5^. While the BL1 motif adopts an open loop conformation outside of the proteasome, it assumes an ordered β-hairpin structure in the proteasome. The BL2 conformation in the proteasome is slightly changed from that in the crystal structure of UbAl-bound USP domain but is moved ~4 Å from that in the ubiquitin-free USP domain structure. **c,** Closeup view of ubiquitin interaction with the BL1 motif and of USP interaction with the OB domains of RPT1-RPT2 in state E_D2_^USP14^. Key residues in BL1, BL2 and BL3 mediating the USP-OB interactions are labelled. **d,** Closeup view of the interface between the USP14 BL1 motif and the RPT OB domain. The residues labelled in red in panels (**c**) and (**d**) correspond to the sites mutated in the USP14 variants shown in panels (**e**) and (**f**). **e,** *In vitro* degradation of Ub_*n*_-Sic1^PY^ by the 26S proteasome assembled with USP14 variants at 37 °C. Samples were analyzed by SDS–PAGE/Western blot using anti-T7 antibody that was used to examinate fusion protein T7-Sic1^PY^. These experiments were repeated independently three times with consistent results. **f**, Ubiquitin–AMC hydrolysis by the USP14 mutants measures their DUB activity in the human proteasome. RFU, relative fluorescence units. Since the RFU increased with time, RFU at 60 min was used to draw the histogram of column. Data are presented as mean ± s.d. from three independent experiments. The quantification on the wildtype USP14 was used as a denominator to normalize all measurements in each experiment. Dots, individual data points.

USP14 interacts with both the OB and AAA domains of the ATPase ring. The USP-OB interaction is mediated by the blocking loops 1 (BL1), 2 (BL2) and 3 (BL3) of the USP domain (Fig. 2b), which buries a solvent-accessible area of approximately 527 Å^2^ on USP14. BL1 makes the most extensive interface with the OB ring around residues Gln128 and Asp133 of RPT1, whereas Ser430 in BL2 and Trp472 in BL3 make brief interactions with Gln128 of RPT1 and Asp118 of RPT2, respectively (Fig. 2c). Between different proteasome states, the movement of the USP domain appears to adapt to the rocking of the OB ring and to maintain its interactions with the OB domains of RPT1-RPT2 (Fig. 3a, Extended Data Fig. 7a).

**Figure 3.**
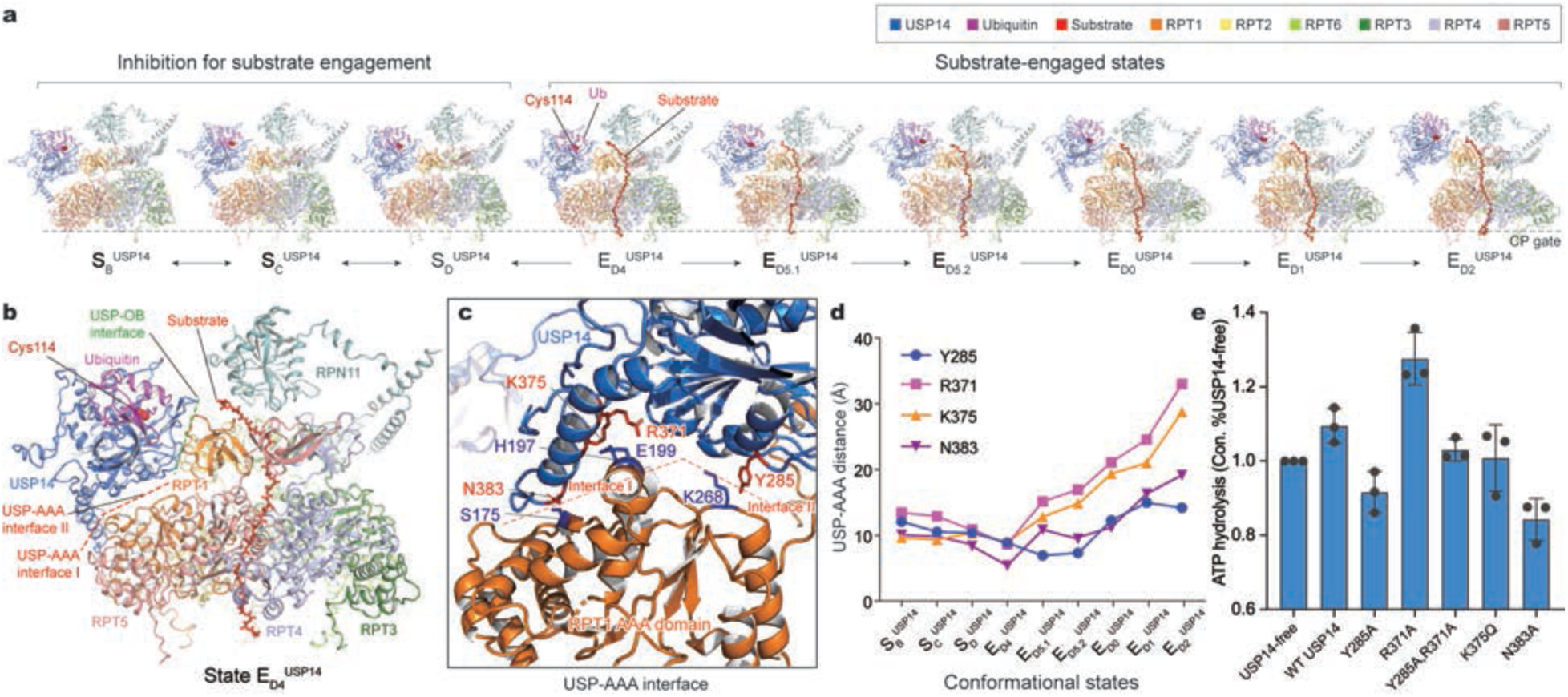
Structural dynamics and mechanism of allosteric regulation of the AAA-ATPase motor by USP14. **a**, Side-by-side comparison of the USP14-ATPase subcomplex structures in six substrate-engaged states (E_D4_^USP14^, E_D5.1_^USP14^, E_D5.2_^USP14^, E_D0_^USP14^, E_D1_^USP14^ and E_D2_^USP14^) and three substrate-inhibited states (S_B_^USP14^, S_B_^USP14^ and S_C_^USP14^). All structures are aligned with respect to their CP components. **b**, Closeup side-view of the USP14-ATPase subcomplex structure in state E_D4_^USP14^ highlights the relationship of the USP-AAA and USP-OB interfaces. **c**, Closeup view zoomed into the USP-AAA interfaces between USP14 and RPT1, with the interacting pairs of residues highlighted in stick representation. The green and red dashed lines in (**b**) and (**c**) mark the USP-OB interface and the USP-AAA interface I (major) and II (minor), respectively. **d**, Changes of the distance between USP domain of USP14 and the AAA domain of RPT1 in nine conformational states of the USP14-bound proteasome. The USP-AAA distance was characterized by measuring the shortest distance between USP14 residues (Y285, R371, K375 and N383) and the main chains of RPT1 AAA domain. **e**, The ATPase activity was quantified using the malachite analysis, by measuring the concentration of phosphate from ATP hydrolysis of the proteasome. Data are presented as mean ± s.d. from three independent experiments. The quantification on the USP14-free proteasome was used as a denominator to normalize all measurements in each experiment. Dots, individual data points.

In contrast to the USP-OB interface that is nearly invariant in all states, USP-AAA interactions vary prominently (Fig. 3a). In states E_D0_^USP14^, E_D1_^USP14^ and E_D2_^USP14^, the USP domain is flipped up and completely detached from the AAA domain of RPT1. By contrast, the USP domain exhibits differential interactions with the AAA domain of RPT1 in other states (Fig. 3b, c). The most extensive USP-AAA interactions are observed in state E_D4_^USP14^ (Fig. 3d). In this conformation, a helix-loop region (residues 371-391) protruding from the USP domain contacts the AAA domain of RPT1, which buries a solvent-accessible area of approximately 850 Å^2^ on USP14 (Fig. 3b, c). The overall USP14 structure bridges the RPN1 and RPT1 subunits in parallel to the RPT1-PRT2 coiled coil domains that directly associate with RPN1. The RPN1-RPT1-RPT2 interface transmits the ubiquitin and UBL interactions on RPN1 to the allosteric regulation of the AAA-ATPase motor^10^. By establishing a parallel structural connection, USP14 modifies how RPN1 allosterically impacts the AAA-ATPase conformation. Indeed, structural comparisons exhibit differential rotations of RPN1 and the lid relative to the base induced by USP14 (Extended Data Fig. 5b).

## USP14 activation by the proteasome

Sequestration of the ubiquitin C-terminus binding groove in USP14 by BL1 and BL2 auto-inhibits the DUB activity in the absence of the proteasome^5^. Comparison with the crystal structures of the USP domain in isolated and ubiquitin aldehyde (UbAl)-bound forms^5^ reveals differential conformational changes in BL1 and BL2 (Fig. 2b). The BL1 region is an open loop in the crystal structures but is refolded into a β-hairpin sandwiched between the OB ring and ubiquitin in the proteasome (Fig. 2b, c). This quaternary architecture suggests that ubiquitin binding stabilizes the BL1 β-hairpin conformation and its interaction with the OB ring, thus rationalizing the previous finding that polyubiquitylated substrates enhance the USP14’s association with the proteasome^18^. In contrast, the BL2 loop is moved approximately 4 Å to make way for ubiquitin C-terminus docking to the groove at the active site. Stabilized by the USP-OB interface, BL1 and BL2 together hold the ubiquitin C-terminus in a β-strand conformation (Fig. 2c), where the mainchain carbon of ubiquitin C-terminal Gly76 is placed approximately 3.4 Å away from the sulfur atom of the catalytic Cys114 and is fully detached from the substrate, indicating that the structure represents a post-deubiquitylation state of USP14.

To test the functional importance of USP-OB interface in USP14 activation in the proteasome, we performed structure-based site-directed mutagenesis using our in vitro-reconstituted degradation system (Extended Data Fig. 8). Indeed, the BL1 mutations, including single mutant K334Q, double mutant V343A/L344A at BL1-OB interface (Fig. 2d) and F331A/Y333A at the ubiquitin-BL1 interface (Fig. 2c), all abrogated the DUB activity of USP14 and showed no obvious inhibition of proteasomal degradation (Fig. 2e, f). By contrast, the single mutants F332A in BL1 and W472A in BL3 still retained a reduced DUB activity and inhibited proteasome function (Fig. 2e, f). These contrasting phenotypes substantiate our structural finding that the BL1-OB interaction is essential for USP14 activation toward efficacious deubiquitylation.

Unexpectedly, the overall structure of the full-length USP14 exhibits three major states (Extended Data Fig. 7b), far less than the number of USP14-bound proteasome states, suggesting the conformational entropy of the linker region is greatly reduced upon USP14 assembly and activation on the proteasome. To test whether the linker length or composition would impact the USP14 activation by the proteasome, we made three mutants with deletion of residues 93-96, insertion of TEEQ after residue 92, and double point-mutations of E90K/D91A. While all three USP14 mutants exhibited inhibition of proteasomal degradation of Ub_*n*_-Sic1^PY^, they also showed 20-30% reduction in the DUB activity relative to the wildtype USP14 (Extended Data Fig. 8c, d). These observations suggest that the linker length and composition in the wildtype USP14 may have been evolutionarily optimized for the DUB activity, although the UBL-USP domain architecture appears functionally robust against the variation of the linker region.

## Allosteric regulation of ATPase activity by USP14

States E_D4_^USP14^, E_D5.1_^USP14^ and E_D5.2_^USP14^ are specifically induced by USP14, since no comparable conformations were reconstructed in the absence of USP14 previously despite exhaustive 3D classification^10, 25^. A common feature of these newly observed substrate-engaged states is a buried surface area of USP-AAA interfaces, suggesting that USP14 preferentially recognizes these AAA conformations and stabilizes these states by direct interactions (Figs. 3a, 4a, Extended Data Fig. 5). In state E_D4_^USP14^ structure, USP14 interacts with the AAA domain of RPT1 via two discrete interfaces (Fig. 3b). The primary USP-AAA interface is mediated by a negatively charged surface on a helix (residues 372-383) and a loop (residues 383-391) protruding out of the USP domain, which buries the majority of the USP-AAA interfaces. This helix-loop region appears to pack against a ridged surface spanning from the large and small AAA subdomains of RPT1, where Arg371, Lys375 and Asn383 of USP14 interact with Ser175, His197 and Glu199 of RPT1. The second USP-AAA interface is centered on Tyr285 in the PKL loop region of USP14 that briefly interacts with Lys267 and Lys268 of RPT1.

**Figure 4.**
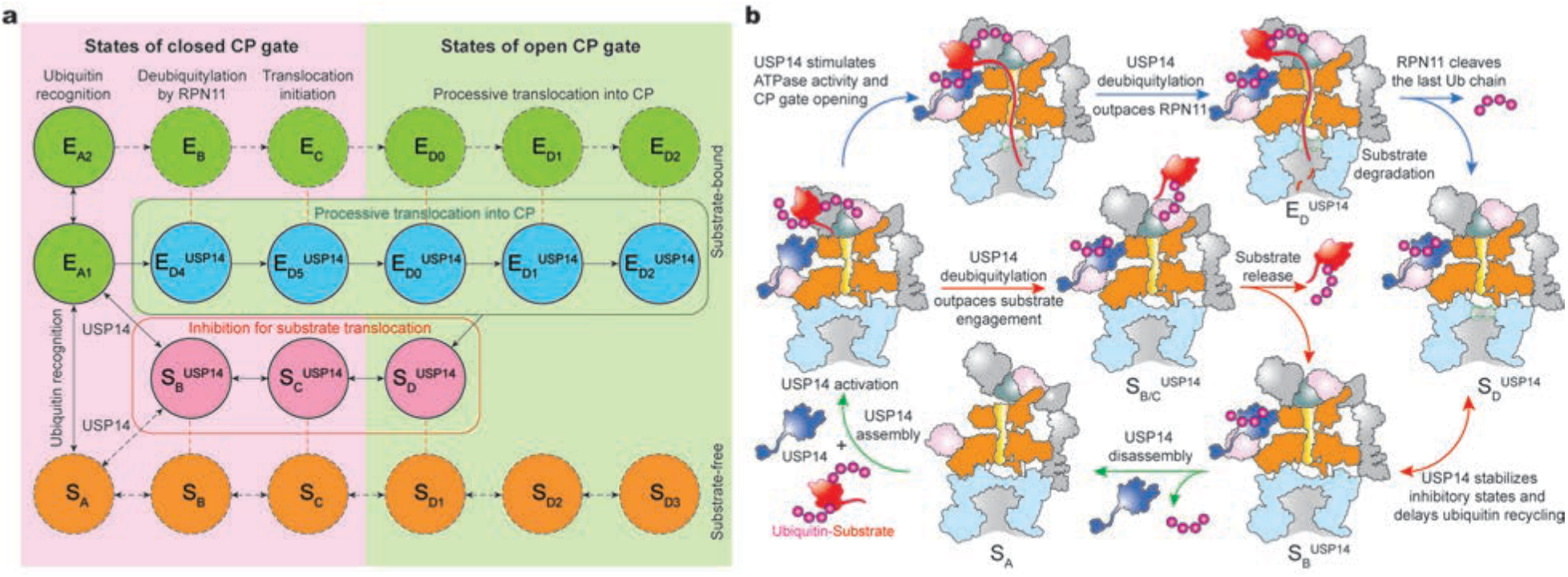
Mechanistic insights into USP14-mediated regulation of proteasome function. **a**, An integrated schematic diagram of proteasome state transitions illustrates the full functional cycles of the proteasome in the presence and absence of USP14. The solid circles are the states observed in the current study, whereas the dashed circles are the states observed in previous studies of substrate-free^8^ (orange) or substrate-engaged human proteasome^10^ (green) in the absence of USP14. Color blue and salmon label the substrate-engaged and substrate-inhibited USP14-proteasome states, respectively. The states with closed and open CP gate are placed in pink and limon backgrounds, respectively. Vertical orange dashed lines link the state pairs with comparable AAA-ATPase structures. Black arrows link the putative pairs of state transitions along a structurally feasible reaction pathway. **b**, Proposed model of USP14 activation-deubiquitylation-disassembly cycle in the proteasome. During the step of USP14 assembly onto the proteasome, ubiquitin-substrate conjugates recruited to the proteasome’s ubiquitin receptor promotes USP14 binding to the RPN1 and RPT1 subunits of the proteasome, which activates USP14. USP14 binding to the proteasome creates two parallel state-transition pathways. The substrate-inhibited pathway, marked by red arrows, has RPN11 blocking the substrate entrance of OB ring before any substrate insertion takes place. Along this pathway, USP14 trims ubiquitin chains and release the substrate from the proteasome, thus preventing the substrate degradation. This pathway is characterized by the cryo-EM structures of states S_B_^USP14^, S_C_^USP14^ and S_D_^USP14^. The substrate-engaged pathway, marked by blue arrows, has RPN11 closed on the OB entrance after a substrate has already inserted into the ATPase ring. As USP14 favors the E_D_-like state of the proteasome, it allosterically stimulates the ATPase activity and promotes the ATPase-CP interaction to open the CP gate early. Along this pathway, USP14 deubiquitylation outpaces that of RPN11 until the last ubiquitin chain remains on a nearby ubiquitin receptor, which is presumably cleaved by RPN11 for the completion of substrate degradation. This pathway is characterized by the cryo-EM structures of states E_D4_^USP14^, E_D5.1_^USP14^, E_D5.2_^USP14^, E_D0_^USP14^, E_D1_^USP14^ and E_D2_^USP14^. Although our data do not intuitively explain why USP14 trim ubiquitin until the last one on a substrate remains, they collectively implicate that the ubiquitin recognition by USP14 in the proteasome likely requires at least another helper ubiquitin chain that is already anchored on a nearby ubiquitin receptor. This ubiquitin chain may not be available for USP14 binding but can be readily trimmed by RPN11.

To test the functional roles of the USP-AAA interfaces, we mutated several USP14 residues at the interfaces to alanine. Although none of these USP14 mutants exhibited any observable defects in the DUB activity and proteasome inhibition (Extended Data Fig. 8e), they demonstrated notable variations in the ATPase activity in the presence of substrate. Consistent with previous studies^16–20^, Ub_*n*_-Sic1^PY^ stimulated the ATPase activity in the wildtype USP14-proteasome by approximately 10% relative to that in the USP14-free proteasome. Two mutants Y285A and N383A showed approximately 20-30% reduction of the ATPase activity to a level 10-20% below that of the USP14-free proteasome (Fig. 3e). As Tyr285 is located halfway between the primary USP-AAA and USP-OB interfaces (Fig. 3c), it may be mechanically important in transmitting the allosteric impact from USP to AAA. By contrast, the single mutant R371A enhanced the ATPase activity by approximately 20% relative to the wildtype USP14 likely via tightening the USP-AAA interactions (Fig. 3c). Intriguingly, the double mutant Y285A/R371A restored the ATPase rate to the approximate level of the USP14-free proteasome, presumably due to the cancelation of two counteracting allosteric effects between the two mutated residues (Fig. 3e). Altogether, our structure-guided mutagenesis validates that the USP-AAA interfaces mediate the non-catalytic, allosteric regulation of the ATPase activity in the proteasome.

## Asymmetric ATP hydrolysis around the AAA-ATPase ring

The six substrate-engaged USP14-proteasome structures characterize a detailed intermediate sequence of substate translocation in the USP14-regulated proteasome, which is substantially deviated from the pathway of translocation initiation in the USP14-free proteasome^10^ (Extended Data Fig. 5). Given the absence of the E_B_- and E_C_-like states in the presence of USP14, these structural observations together prompt the hypothesis that states E_D4_^USP14^ and E_D5_^USP14^ may replace the USP14-free states EB and E_C_ during initiation of substrate translocation in the USP14-bound proteasome. Thus, we infer that these states present a continuum of USP14-altered conformations following the state-transition sequence of E_A1_→ E_D4_^USP14^→ E_D5.1_^USP14^→ E_D5.2_^USP14^→ E_D0_^USP14^→ E_D1_^USP14^→ E_D2_^USP14^ (Fig. 4a). In support of this sequence assignment, the RPT3 pore-1 loop is moved sequentially from the top to the bottom of the substrate-pore loop staircase, whereas coordinated ATP hydrolysis navigates around the ring for a near-complete cycle (Extended Data Fig. 5b, e). A hypothetical state E_D3_^USP14^ connecting E_D2_^USP14^ and E_D4_^USP14^ was not yet observed.

The newly resolved states fill up a major missing gap in our current understanding of substrate translocation in the proteasomal AAA-ATPase. It has remained unclear whether a strict sequential hand-over-hand mechanism is used by the AAA-ATPase for processive substrate translocation^26^. To our surprise, four of the six E_D_-compatible states—E_D5.1_^USP14^, E_D5.2_^USP14^, E_D0_^USP14^ and E_D1_^USP14^— exhibit an AAA-ATPase structure with two adjacent RPT pore-1 loops disengaged from the substrate: one moving away from the substrate in the process of releasing ADP for nucleotide exchange, and the other moving up toward the substrate in the process of binding ATP for substrate re-engagement (Extended Data Figs. 5, 9). Recent studies have found that the substrate re-engagement of an ATPase subunit can be the rate-limiting step in single-nucleotide exchange dynamics^25^. Coupling of conformational changes between adjacent AAA domains may vary from tightly coupled kinetics—i.e., only one dissociated pore-1 loop at a time—to loosely coupled kinetics exhibiting two adjacent pore-1 loops dissociated from substrate simultaneously^10, 25^. Thus, two adjacent pore-1 loops dissociated from the substrate represent an inevitable intermediate in the sequential hand-over-hand model of substrate translocation, which would be very short-lived in tightly coupled kinetics^27^. Under the allosteric influence of USP14, such kinetics can become loosely coupled around RPT1, leading to meta-stabilization of these short-lived intermediate states. The observations of both tightly and loosely coupled kinetics at different locations around the ATPase ring rationalize broken symmetry of coordinated ATP hydrolysis in the ATPase motor—an effect that has been previously informed by functional studies^28, 29^.

## Insights into USP14-mediated proteasome regulation

Our structural, kinetic and functional data collectively provide novel insights into how USP14 regulates the proteasome activity at multiple checkpoints by inducing parallel pathways of proteasome state transitions (Fig. 4a). The first checkpoint is at the initial ubiquitin recognition prior to substrate engagement with the ATPases. USP14 assembly on the proteasome is facilitated by ubiquitin binding on USP14 that appears to stabilize several substrate-inhibited states such as S_B_^USP14^ and S_C_^USP14^ (Fig. 4b). Should ubiquitin be recruited to RPN11, USP14 binding is then mutually excluded, as observed in state E_A2_. Such a competition in ubiquitin recruitment coerces the proteasome to bypass the conformational transition pathway represented by USP14-free states E_A2_, E_B_ and E_C_, which are needed to couple RPN11-catalysed deubiquitylation with ATP-dependent substrate engagement and translocation initiation^10^.

However, the stabilization of substrate-inhibited states does not completely exclude the likelihood of substrate engagement with the ATPases. Substrate insertion into the ATPase ring can stochastically occur before USP14 activation by the proteasome. In the presence of ubiquitylated substrates, USP14 stimulates the ATPase rate^16–18^ and induces early CP gate opening, represented in state E_D4_^USP14^, E_D5.1_^USP14^ and E_D5.2_^USP14^. Thus, USP14 creates a second kinetic checkpoint and drives the proteasome to choose between two alternative pathways of USP14-regulated conformational transitions—one that sterically prevents the substrate engagement and the other that decouples the DUB activity of RPN11 from substrate engagement and translocation. In support of this model, all substrate-engaged USP14-proteasome states showed no ubiquitin binding to RPN11.

Catalytically, USP14 trims ubiquitin chains from the substrate quite fast^7^. For this reason, we have not captured the USP14 conformation right at the moment of deubiquitylation. Upon completion of substrate degradation, the substrate-engaged states of USP14-proteasomes are partly converted into substrate-inhibited states (Fig. 1f). Given that all observed states also present USP14-bound ubiquitin, the lack of observation on ubiquitin-free USP14 in the proteasome implicates that USP14 delays the release of trimmed ubiquitin chains from the proteasome, since the USP14-bound ubiquitin stabilizes the USP-OB interaction (Fig. 2c). Thus, it is conceivable that ubiquitin binding is needed for USP14 activation on the proteasome and the cleaved ubiquitin chain is released upon the USP14 disassembly from the proteasome^16, 17^ (Fig. 4b), in contrast to RPN11 that recycles cleaved ubiquitin quickly^10^. This creates a third checkpoint, where the polyubiquitin-bound USP14 kinetically delays ubiquitin recycling from the proteasome and slows down additional substrate recruitment (Fig. 4b).

In summary, USP14 acts as an adaptive regulator of the AAA-ATPase motor and parallels the RPN1-RPT1-RPT2 regulatory axis in a switchable fashion. USP14 interactions kinetically create three branching ‘decision-making’ checkpoints on the proteasome, at the steps of initial ubiquitin recognition, of substrate translocation initiation by the AAA-ATPase motor, and of recycling trimmed ubiquitin chains. While the first checkpoint allows USP14 to allosterically prevent RPN11 from accepting substrate-conjugated ubiquitin, the second checkpoint either directly rejects substrate commitment or kinetically antagonizes RPN11 by outpacing the coupling of RPN11-directed deubiquitylation and substrate-ATPase engagement, rather than directly inhibiting the RPN11’s DUB activity. This is achieved at the cost of stimulating the ATPase activity and of tightening the RP-CP interface. The third checkpoint further controls the ubiquitin-recycling function of the proteasome that is critical for regulating the free ubiquitin reservoir in cells. This multi-checkpoint mechanism integrates catalytic and non-catalytic effects of proteasome inhibition into a comprehensive, elegantly regulated process of substrate degradation. As partly supported by another study on the yeast Ubp6-bound proteasome^30^, such a mechanism is expected to be conserved from yeast to human and shed light on how reversibly associated DUBs and other proteins regulate the proteasome function in general.

## Methods

### Expression and purification of human USP14

Wild-type USP14 and mutants were cloned into pGEX-4T vector obtained from GenScript (Nanjing, China). For purification of recombinant USP14 and mutants, BL21-CondonPlus (DE3)-RIPL cells (Shanghai Weidi, China) transformed with plasmids encoding wild-type or mutant USP14 were grown to an OD_600_ of 0.6-0.7 in LB medium supplemented with 100 mg/ml ampicillin. Cultures were cooled to 20 °C and induced with 0.2 mM IPTG overnight. Cells were harvested by centrifugation at 3000 × g for 15 min and resuspended in lysis buffer (25 mM Tris-HCl [pH 8.0], 150 mM NaCl, 0.2% NP-40, 1 mM DTT, 10% glycerol and 1 × protease inhibitor cocktail). Cells were lysed by sonication and the lysate was cleared through centrifugation at 20000 × g for 30 min at 4 °C. The supernatant was incubated with glutathione Sepharose 4B resin (GE Healthcare) for 3 h at 4 °C. For wild-type USP14 purification, the resin was washed with 20 column volumes of washing buffer (25 mM Tris-HCl [pH 8.0], 150 mM NaCl, 1 mM DTT, 10% glycerol), then incubated with cleavage buffer (20 mM Tris-HCl [pH 8.0], 150 mM NaCl) containing thrombin (Sigma) overnight at 4 °C. The eluted samples were further purified by gel-filtration column (Superdex 75, GE Healthcare) equilibrated with 25 mM Tris-HCl [pH 8.0], 150 mM NaCl, 1 mM DTT, 10% glycerol. For the purification of USP14 mutants, the resin was washed with 20 column volumes of washing buffer (25 mM Tris-HCl [pH 8.0], 300 mM NaCl, 1 mM DTT), then incubated with cleavage buffer (20 mM Tris-HCl [pH 8.0], 150 mM NaCl) containing thrombin (Sigma) overnight at 4 °C. To remove thrombin, the GST eluent was incubated with Benzamidine-Sepharose (GE Healthcare) for 30 min at 4 °C.

### Expression and purification of human 26S proteasome

HTBH (hexahistidine, TEV cleavage site, biotin, and hexahistidine)-tagged human 26S proteasomes were affinity purified from a stable HEK293 cell line (a gift from L. Huang, University of California, Irvine) as previously described^8–10^. For ubiquitin-vinyl-sulfone (Ub-VS) treated human proteasome, 1 μM Ub-VS (Boston Biochem) was added to the proteasomes binding resin and incubated for 2 h at 30 °C. Residual Ub-VS was removed by washing the beads with 30 bed volumes of wash buffer (50 mM Tris-HCl [pH7.5], 1 mM MgCl_2_ and 1 mM ATP). The proteasomes were cleaved from the beads using TEV protease (Invitrogen) and used to measure the DUB activity of USP14 using the Ub–AMC hydrolysis assay.

### Preparation of polyubiquitylated Sic1^PY^

Sic1^PY^ and WW-HECT were purified as previously described^10^. Human UBE1 (plasmid obtained as a gift from C. Tang, Peking University) and human UBCH5A (obtained from GenScript, China) were expressed as GST fusion proteins from pGEX-4T vectors. In brief, UBE1-expressing BL21-CondonPlus (DE3)-RIPL cell cultures were induced with 0.2 mM IPTG for 20 h at 16°C, whereas UBCH5A expression was induced with 0.2 mM IPTG overnight at 18°C. Cells were harvested in lysis buffer (25 mM Tris-HCl [pH 7.5], 150 mM NaCl, 10 mM MgCl_2_, 0.2% Triton-X-100, 1 mM DTT) containing 1× protease inhibitor cocktail and lysed by sonication. The clear lysates were incubated with glutathione Sepharose 4B resin for 3 h at 4 °C and subsequently washed with 20 bed volumes lysis buffer. The GST tag was removed by thrombin protease (Sigma) in cleavage buffer (20 mM Tris-HCl [pH 8.0], 150 mM NaCl, 1mM DTT) overnight at 4°C. The eluted samples were further purified by gel-filtration column (Superdex 75, GE Healthcare) equilibrated with 25 mM Tris-HCl [pH 7.5], 150 mM NaCl, 1 mM DTT, 10% glycerol.

To ubiquitylate Sic1^PY^, 1.2 μM Sic1^PY^, 0.5 μM UBE1, 2 μM UBCH5a, 1.4 μM WW-HECT and 1 mg/ml ubiquitin (Boston Biochem) were incubated in reaction buffer (50 mM Tris-HCl [pH 7.5], 100 mM NaCl, 10 mM MgCl_2_, 2 mM ATP, 1 mM DTT and 10% glycerol) for 2 h at room temperature. His-tagged Sic1^PY^ conjugates (polyubiquitylated Sic1^PY^, Ub_*n*_-Sic1^PY^) were purified by incubating with Ni-NTA resin (Qiagen) at 4 °C for 1 h. After the resin was washed with 20 column volumes of the wash buffer (50 mM Tris-HCl [pH 7.5], 100 mM NaCl, 10% glycerol), the Ub_*n*_-Sic1^PY^ was eluted with the same buffer containing 150 mM imidazole, and finally exchanged to the storage buffer (50 mM Tris-HCl [pH 7.5], 100 mM NaCl, 10% glycerol) using an Amicon ultrafiltration device with 30K molecular cut-off (Millipore).

### Expression and purification of human RPN13

To purify the human RPN13, pGEX-4T-RPN13-transformed BL21-CondonPlus (DE3)-RIPL cells were cultured to an OD_600_ of 0.6 and then induced by 0.2 mM IPTG for 20 h at 16 °C. Cells were resuspended in lysis buffer (25 mM Tris-HCl [pH 7.5], 300 mM NaCl, 1 mM EDTA, 0.2% Triton-X-100, 1 mM DTT) containing 1× protease inhibitor cocktail and lysed by sonication. A 20,000 × g supernatant was incubated with glutathione Sepharose 4B resin (GE Healthcare) for 3 h at 4 °C. The resin was washed with 20 column volumes of washing buffer (25 mM Tris-HCl [pH 8.0], 150 mM NaCl, 1 mM DTT, 10% glycerol) and 10 column volumes of cleavage buffer (20 mM Tris-HCl [pH 8.0], 150 mM NaCl). GST tag was cleaved by incubating with thrombin (Sigma) overnight at 4°C. The eluted samples were further purified by gel-filtration column (Superdex 75, GE Healthcare) equilibrated with 25 mM Tris-HCl (pH 8.0), 150 mM NaCl, 1 mM DTT, 10% glycerol.

### *In vitro* degradation assay

Purified human proteasomes (~30 nM) were incubated with RPN13 (~300 nM), Ub_*n*_-Sic1^PY^ (~300 nM) in degradation buffer (50 mM Tris-HCl [pH 7.5], 5 mM MgCl_2_ and 5 mM ATP) at 37 °C. Purified recombinant USP14 variants (~1.2 μM) were incubated with proteasome for 20 min at room temperature before initiating the degradation reaction. The reaction mixtures were incubated at 37 °C for 0, 0.5, 1.0 and 2.0 min, or 10 °C for 0, 2.0, 5.0 and 10 min, then terminated by adding SDS loading buffer and subsequently subjected to western blot using anti-T7 antibody (Abcam) that was used to examinate fusion protein T7-Sic1^PY^.

### Ubiquitin-AMC hydrolysis assay

Ubiquitin-AMC (Ub-AMC; Boston Biochem) hydrolysis assay was used to verify the wild-type and mutant USP14 deubiquitylating activities in the human proteasome. The reactions were performed in reaction buffer (50 mM Tris-HCl [pH 7.5], 5 mM MgCl_2_, 1 mM ATP, 1 mM DTT, 1 mM EDTA and 1 mg/mL ovalbumin [Diamond]), containing 1 nM Ub-VS-treated proteasome, 0.2 μM USP14 variants and 10 nM RPN13. The reaction was initiated by adding 1 μM Ub-AMC. Ub-AMC hydrolysis was measured in a Varioskan Flash spectral scanning multimode reader (Thermo Fisher) by monitoring an increase of fluorescence excitation at 345 nm with an emission at 445 nm.

### ATPase activity assay

ATPase activity was quantified using the malachite green phosphate assay kits (Sigma). Human proteasomes (30 nM), RPN13 (300 nM) and USP14 variants (1.2 μM) were incubated in assembly buffer (50 mM Tris-HCl [pH 7.5], 5 mM MgCl_2_ and 5 mM ATP) for 20 min at room temperature, subsequently added with Ub_*n*_-Sic1^PY^ (300 nM) and incubated for 1 min at 37°C. The reaction mixtures were mixed with malachite green buffers as described by the manufacturer (Sigma). After 30 min of room temperature incubation, the O.D. at 620 nm was determined using a Varioskan Flash spectral scanning multimode reader (Thermo Fisher).

### Cryo-EM sample preparation

To prepare cryo-EM samples, all purified proteins were exchanged to imaging buffer (50 mM Tris-HCl [pH 7.5], 5 mM MgCl_2_ and 1 mM ATP). 1 μM human proteasomes were incubated with 10 μM RPN13, 10 μM USP14 in imaging buffer (50 mM Tris-HCl [pH 7.5], 5 mM MgCl_2_ and 1 mM ATP) for 20 min at 30 °C, then cooled to 10 °C. 10 μM Ub_*n*_-Sic1^PY^ was added to the mixture and incubated at 10 °C for 0, 1, 5 and 10 min. 0.005% NP-40 was added to the reaction mixture immediately before cryo-plunging. For ATP-to-ATPγS exchange and ATPase quenching, after the reaction mixture was incubated at 10 °C for 1 min, 1 mM ATPγS was added to the reaction mixture at once, and incubated for another 1 min, then NP-40 was added to the mixture to a final concentration of 0.005% before cryo-plunging.

### Cryo-EM data collection

The cryo-grids were initially screened on a 200 kV Tecnai Arctica microscope (Thermo Fisher). Good-quality grids were then transferred to a 300 kV Titan Krios G2 microscope (Thermo Fisher) equipped with the post-column BioQuantum energy filter (Gatan) connected to K2 Summit direct electron detector (Gatan). Coma-free alignment and parallel illumination were manually optimized prior to each data collection session. Cryo-EM data was acquired automatically using SerialEM software^31^ in a super-resolution counting mode, with the defocus set in the range of −0.8 to −2.0 μm. A total exposure time of 10 s with 250 ms per frame resulted in a 40-frame movie per exposure with an accumulated dose of ~50 electrons/Å^2^. The calibrated physical pixel size and the super-resolution pixel size were 1.37 Å and 0.685 Å, respectively. For time-resolved sample conditions, 1,781, 13,868, 2,073 and 2,071 movies were collected for the reaction time of 0 min, 1 min, 5 min, and 10 min, respectively. For the condition in the presence of ATPγS, 12,210 movies were collected.

### Cryo-EM data processing

Drift correction and dose weighting were performed using the MotionCor2 program^32^ at a super-resolution pixel size of 0.685 Å. Drift-corrected micrographs were used for the determination of the micrograph CTF parameters with the Gctf program^33^. Particles were automatically picked using an improved version of the DeepEM program^34^ and then down-sampled by two-fold to a pixel size of 1.37 Å for the following processing. Micrographs screening and auto-picked particles checking were both preformed in the EMAN2 software^35^. Reference-free 2D classification and 3D classification were carried out in software packages RELION^36^ version 3.1 and ROME^37^. Focused 3D classification, CTF and aberration refinement, and high-resolution auto-refinement were mainly done with RELION 3.1, while the AlphaCryo4D software^38^ was used to analyze the conformational changes and guide the 3D classification. We applied a hierarchical 3D classification strategy to analyze the data (Extended Data Fig. 2), which was optimized according to the method described in a previous paper^10^. The entire data-processing procedure consisted of five steps. Datasets of different conditions were processed separately at step 1, 2 and combined at step 3-5.

#### Step 1

Doubly capped proteasome particles were separated from singly capped ones through several rounds of 2D and 3D classification. These particles were aligned to the consensus models of the doubly and singly capped proteasome to obtain their approximate shift and angular parameters. With these parameters, each doubly capped particle was split into two pseudo-singly capped particles by re-centering the box onto the RP–CP subcomplex. Then the box sizes of pseudo-singly capped particles and true singly capped particles were both shrunk to 640 × 640 pixels. A total of 2,549,617 particles in all datasets were obtained after this step.

#### Step 2

Particles were aligned to the CP subcomplex through auto-refinement, followed by one round of the CTF refinement to correct optical aberration (up to the fourth order), magnification anisotropy, and per-particle defocus together with per-particle astigmatism. After another run of the CP-masked auto-refinement, an alignment-skipped RP-masked 3D classification was performed to omit broken 3D classes and separate the S_A_-like states from the S_D_-like states. The RP subcomplex of the S_D_-like states rotated by a large angle compared to the S_A_-like states, and only in S_D_-like states was the catalytic domain of USP14 observed to bind the OB ring in the proteasome. There were totally 1,401,326 particles in S_A_-like states and 967,196 particles in S_D_-like states in all datasets after this step.

#### Step 3

The S_D_-like particles from different datasets were combined for further analysis except the condition of 0 min reaction time, in which the substrate was not yet added into the reaction system. CP-masked auto-refinement was performed to the S_D_-like particles, followed with two rounds of CTF refinement and another run of CP-masked auto-refinement. Alignment-skipped RP-masked 3D classification was then performed, while conformation changes were analyzed using AlphaCryo4D^38^, which yielded three clusters, designated S_B_-like, S_D_-like, and E_D_-like states. These names were referred to the similar states in the previously published papers^9, 10^.

#### Step 4

Particles in different clusters were individually auto-refined with CP masked. The CP density was then subtracted and the particle box was recentered to the RP subcomplex and shrunk to 240 × 240 pixels. For each cluster, the CP-subtracted particles were subjected to several rounds of RP-masked auto-refinement, alignment-skipped RP-masked 3D classification via AlphaCryo4D analysis^38^, finally resulting in nine major conformational states of the USP14-bound proteasome, named S_B_^USP14^, S_C_^USP14^, S_D_^USP14^, E_D0_^USP14^, E_D1_^USP14^, E_D2_^USP14^, E_D4_^USP14^, E_D5.1_^USP14^ and E_D5.2_^USP14^, respectively. The above procedure (step 3 and 4) was also done for the S_A_-like particles, resulting in two major states E_A1_ and E_A2_, which show no clear USP14 binding the OB ring of the proteasome. Time-resolved conformational changes analysis and comparison in the presence and absence of ATPγS could be done after this step, by simply separating the particles for each state based on their time labels.

#### Step 5

After another round of RP-masked 3D classification, incomplete 3D classes (usually with blurred RPN2 or RPN5) were discarded and good-quality particles were selected for the final reconstruction. The final auto-refinement of the CP subcomplex was performed using the pseudo-singly capped particles with a CP mask, while the RP refinement was performed using the CP-subtracted particles. An RP mask and an RPT-RPN1-USP14 mask were separately applied for the RP refinement and the resulting maps were merged into one final map per state. The Fourier shell correlation (FSC) curves of nine states were calculated from two separately refined half maps in a gold-standard procedure, yielding the nominal resolution ranging from 3.0 to 3.6 Å, and the local RP resolution ranging from 3.6 to 6.2 Å. To get a subclass with the USP domain of USP14 tightly bound to the proteasome, a round of USP-masked 3D classification was performed. For the E_D2_^USP14^ state, an USP-tightly-bound RP map at the resolution of 3.8 Å was finally refined, accounting for 40% of the E_D2_^USP14^ particles. Prior to visualization, all density maps were sharpened by applying a negative B-factor calculated by Phenix autosharpen program^39^. Local resolution variations were estimated using ResMap^40^.

### Atomic model building and refinement

Atomic model building was based on the previously published cryo-EM structures of the human proteasome^10^. For the CP subcomplex, initial models of the closed-gate CP and open-gate CP were respectively derived from the E_A1_ model (PDB 6MSB) and the E_D2_ model (PDB 6MSK). For the RP subcomplex, the previous E_D2_ model was used as a template. All subunits of the template model were individually fitted as a rigid body into each of the reconstructed maps with UCSF Chimera^41^, followed by further adjustment of the mainchain traces using Coot^42^. Initial model of the USP14 was derived from a predicted one by AlphaFold^43^, which was verified by comparing to a crystal structure^5^ (PDB 2AYO). The USP14 UBL domain was fitted as a rigid body into the maps, and the linker between the UBL and USP domains was manually fitted in Coot^42^. N-terminus of the subunit RPN5 into the densities of low local resolution was rebuilt by considering the prediction of AlphaFold^43^. Subunit SEM1 was also remodeled by adding residues one by one from the C-terminus in Coot^42^. For some structures with partially blurred or missing subunits density, the atomic models were revised by removing these regions, e.g., the N-termini of both RPN1 and RPN2 of the E_D4_^USP14^ model were removed, and the UBL domain of USP14 was not included in the models of E_D5.1_^USP14^ and E_D5.2_^USP14^. Given that the substrates were not stalled in a homogeneous location during their degradation and that substrate translocation through the proteasome is not sequence-specific, the substrate densities were modelled using polypeptide chains without assignment of amino acid sequence. For states E_D0_^USP14^, E_D1_^USP14^, E_D2_^USP14^, and E_D4_^USP14^, the nucleotide densities are of sufficient quality for differentiating ADP from ATP, which allowed us to build the atomic models of ADP and ATP into their densities (Extended Data Fig. 9c). For other states with the local RP resolution worse than 4.0 Å, the nucleotide types or states were hypothetically inferred based on the densities, the openness of corresponding nucleotide-binding pockets as well as their homologous structural models of higher resolution if available.

After manually atomic building, atomic models were all subjected to the real-space refinement program in Phenix^39^. Both simulated annealing and global minimization were applied with NCS, rotamer and Ramachandran constraints. Partial rebuilding, model correction and density-fitting improvement in Coot^42^ were then iterated after each round of atomic model refinement in Phenix^39^. The refinement and rebuilding cycle were often repeated for three rounds or until the model quality reached expectation (Extended Data Table 1). All figures of structures were plotted in ChimeraX^44^, PyMOL^45^, or Coot^42^. Structural alignment and comparison were performed in both PyMOL^45^ and ChimeraX^44^.

## Data availability

The three-dimensional cryo-EM density maps of all nine USP14-bound proteasome states are deposited into the Electron Microscopy Data Bank (EMDB) (www.emdatasource.org) (under accession numbers to be provided upon formal publication of this manuscript). The corresponding coordinates are deposited in the Protein Data Bank (PDB) (www.wwpdb.org) (with accession numbers to be provided upon formal publication of this manuscript).

## Acknowledgments

The authors thank Y. Saeki for the plasmids expressing Sic1^PY^ and WWHECT; L. Huang for the proteasome-expressing cell lines; C. Tang for the plasmid expressing UBE1; X. Li, Y. Ma, C. Fan for technical support; A. Goldberg for constructive discussions and comments. This work was funded in part by Beijing Natural Science Foundation (grant No. Z180016/Z18J008) and the National Natural Science Foundation of China (grant No. 11774012). The cryo-EM data were collected at the Cryo-EM Core Facility Platform and Laboratory of Electron Microscopy at Peking University. The data processing was performed in the High-Performance Computing Platform at Peking University.

## Author contributions

Y.M. conceived and supervised this study. S.Zou purified the protein complexes, prepared the cryo-EM samples and conducted the biochemical experiments and mutagenesis study. S.Zhang, S.Zou, and D.Y. collected cryo-EM data, analyzed the experimental cryo-EM datasets. S.Zhang refined the density maps and built and refined the atomic models. F.D. and Z.W. contributed to the analysis of the data. Y.M. analyzed the structures and wrote the manuscript with inputs from all authors.

## Correspondence and requests for materials

should be addressed to Y.M.

---

**Extended Data Fig. 1.**
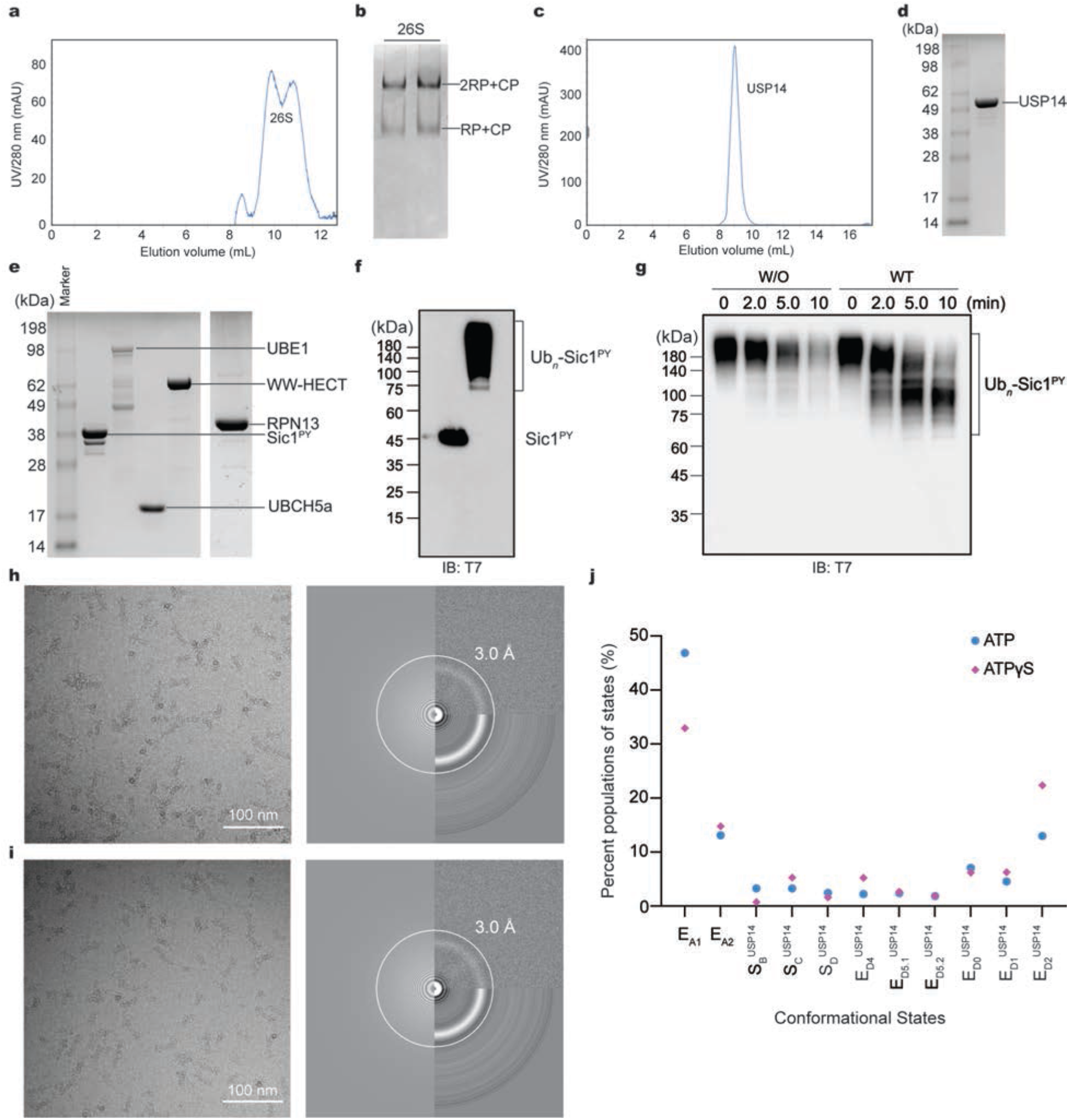
Protein purification and cryo-EM imaging. **a,** The human 26S proteasome was purified through gel-filtration column (Superose 6 10/300 GL). **b,** Native gel analysis of the human 26S proteasome from (**a**). **c,** FPLC purification of human USP14 on Superdex 75 10/300 GL column. **d-c,** SDS-PAGE and Coomassie blue stain analysis of purified USP14, Sic1^PY^, UBE1, UBCH5a, WW-HECT and human RPN13. **f,** Western blot was used to verify polyubiquitylation of Sic1^PY^ (Ub_*n*_-Sic1^PY^) using anti-T7 antibody, indicating almost all Sic1^PY^ was ubiquitylated. **g,** *In vitro* degradation of Ub_*n*_-Sic1^PY^ by the purified 26S proteasome at 10 °C in the absence and presence of USP14. The experiments were repeated 3 times. Samples in (**f**) and (**g**) were analyzed by SDS–PAGE/Western blot using anti-T7 antibody. W/O, the proteasome without binding to USP14. WT, the wildtype USP14-bound proteasome. **h-i,** Typical motion-corrected cryo-EM micrographs (left) of the substrate-engaged human USP14-proteasome complex in the presence of ATP (**h**) or after ATP-to-ATPγS exchange (**i**). Power spectrum evaluation of the corresponding micrographs are shown on the right. **j**, Comparison of percent population of each conformational state in the presence of ATP or ATP-to-ATPγS exchange in 1 min after mixing the substrate with the USP14-bound proteasome.

**Extended Data Fig. 2.**
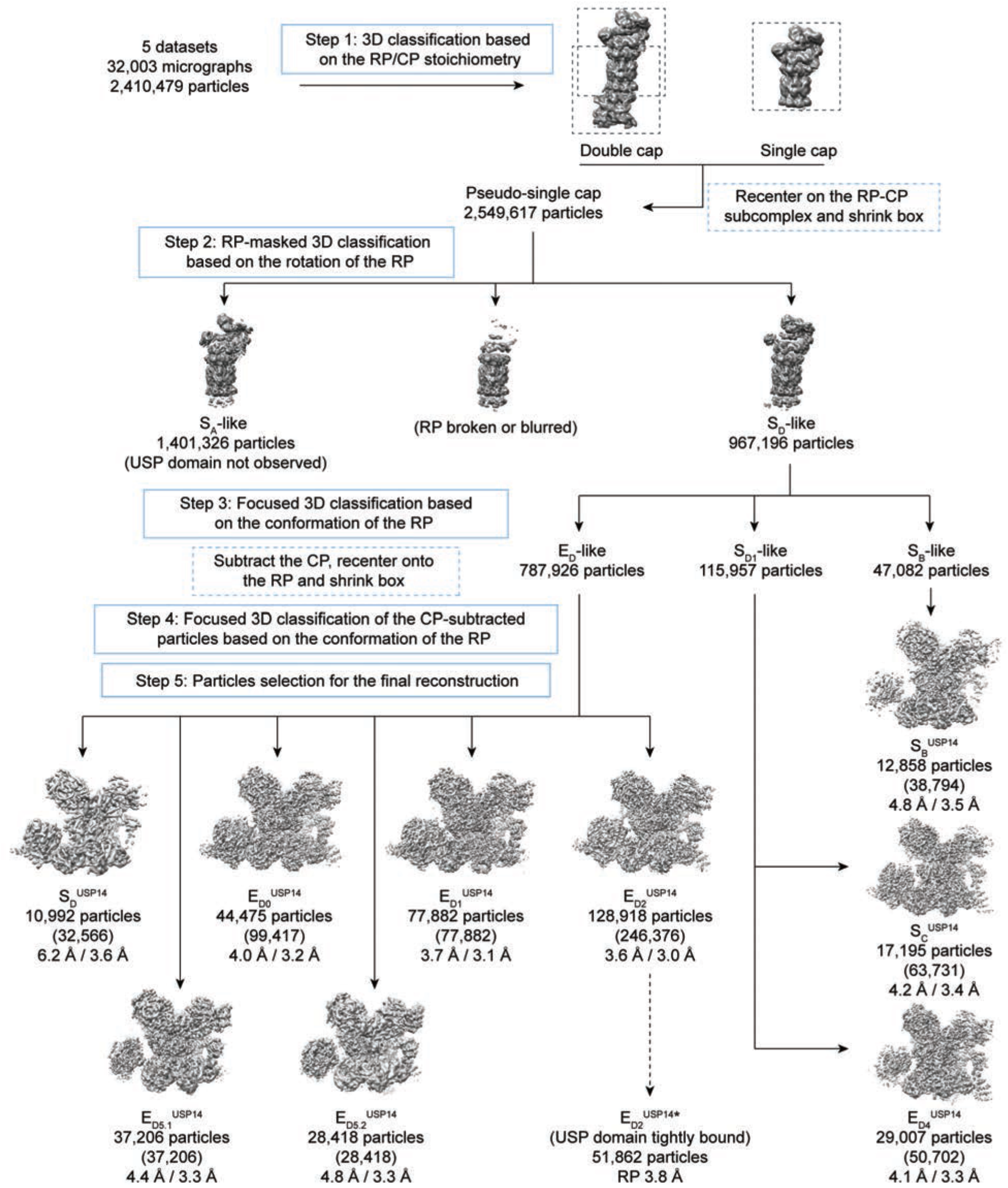
Cryo-EM data processing workflow. The diagram illustrates the five major steps of our focused 3D classification strategy. States E_A1_ and E_A2_, the USP domains of which are invisible, are omitted for clarity. Particles numbers after 3D classification and final reconstruction and the resolution of the RP/CP reconstruction of each state are labelled.

**Extended Data Fig. 3.**
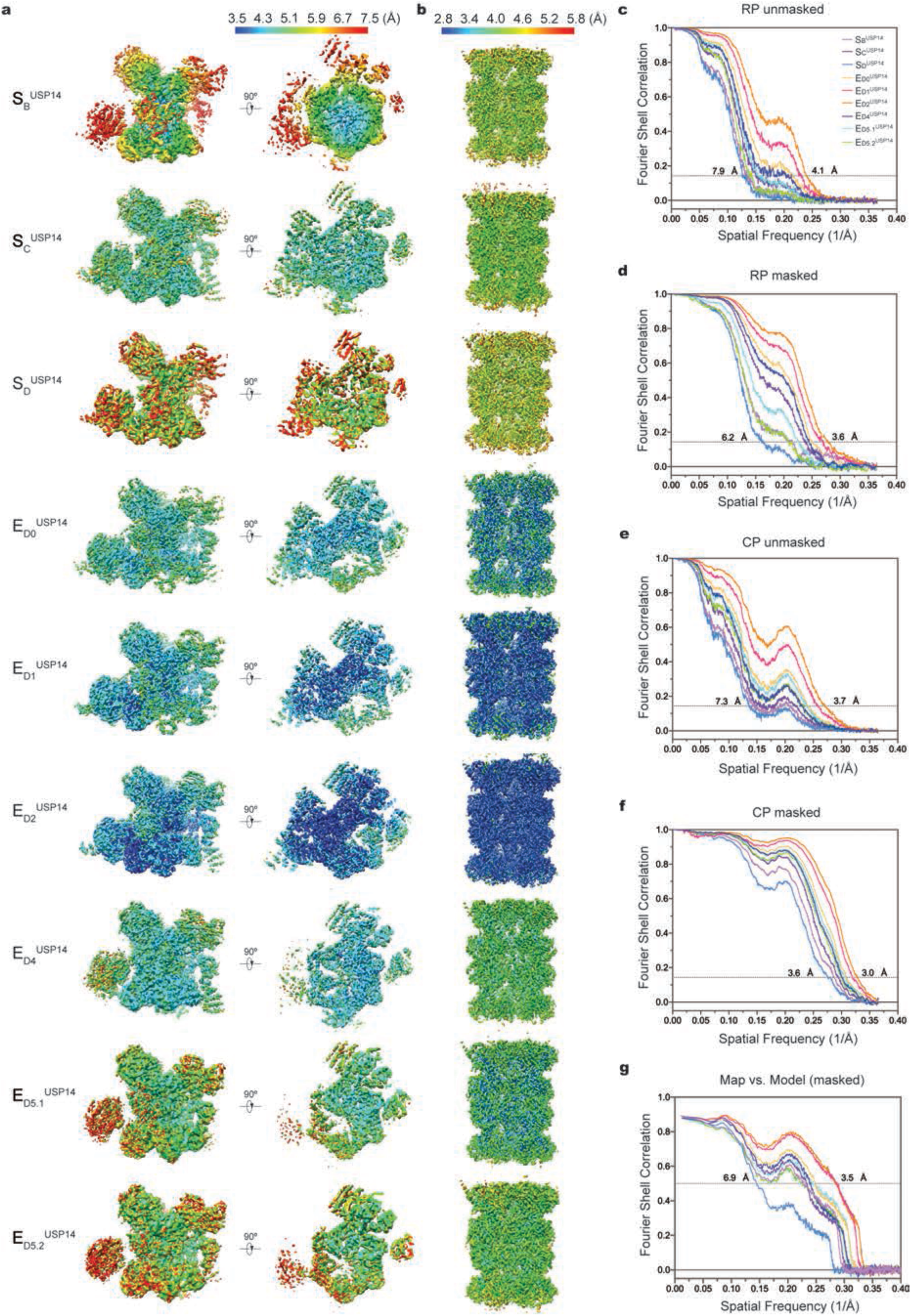
Cryo-EM reconstructions and resolution measurements. **a**, Local resolution estimation of the RP reconstructions of nine states calculated by ResMap^40^. The RP maps were reconstructed using the CP-subtracted particles and shown on the full volumes (left) and on the cross-sections of the AAA ring (right). **b**, Local resolution estimation of the CP reconstructions of nine states. The CP maps were refined by focusing the mask on the CP subcomplex. **c, d**, Gold-standard FSC curves of the RP maps calculated without (**c**) or with (**d**) masking the separately refined half-maps. **e, f**, Gold-standard FSC curves of the CP maps calculated without (**e**) or with (**f**) masking the separately refined half-maps. **g**, Model-map FSC plots calculated by Phenix^39^ between each map (masked) and its corresponding atomic model. For each state, separately reconstructed RP and CP maps were merged in Fourier space into a single map, which was used for the model-map FSC calculation. The same color code is used in (**c-g**).

**Extended Data Fig. 4.**
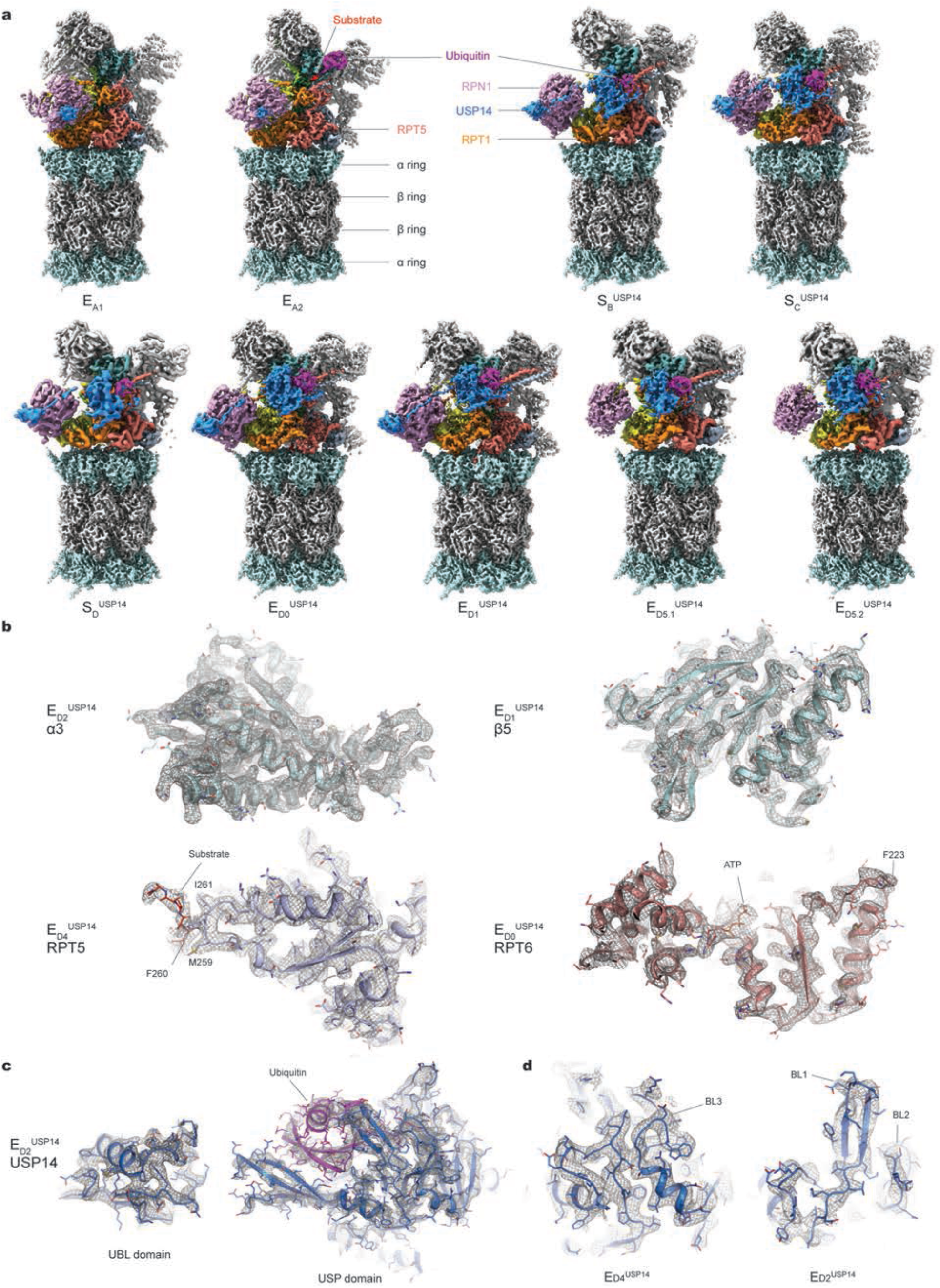
Cryo-EM maps and quality assessment. **a**, Gallery of all refined cryo-EM maps not shown in the main figures. **b-d**, Typical high-resolution cryo-EM densities (grey mesh) of secondary structures superimposed with their atomic models. Different subunits of the proteasome are shown in (**b**), where the substrates shown in the bottom two panels are modelled using polypeptide chains without assignment of amino acid sequence. Two domains of the USP14 and close-up views of its blocking loops are respectively shown in (**c**) and (**d**).

**Extended Data Fig 5.**
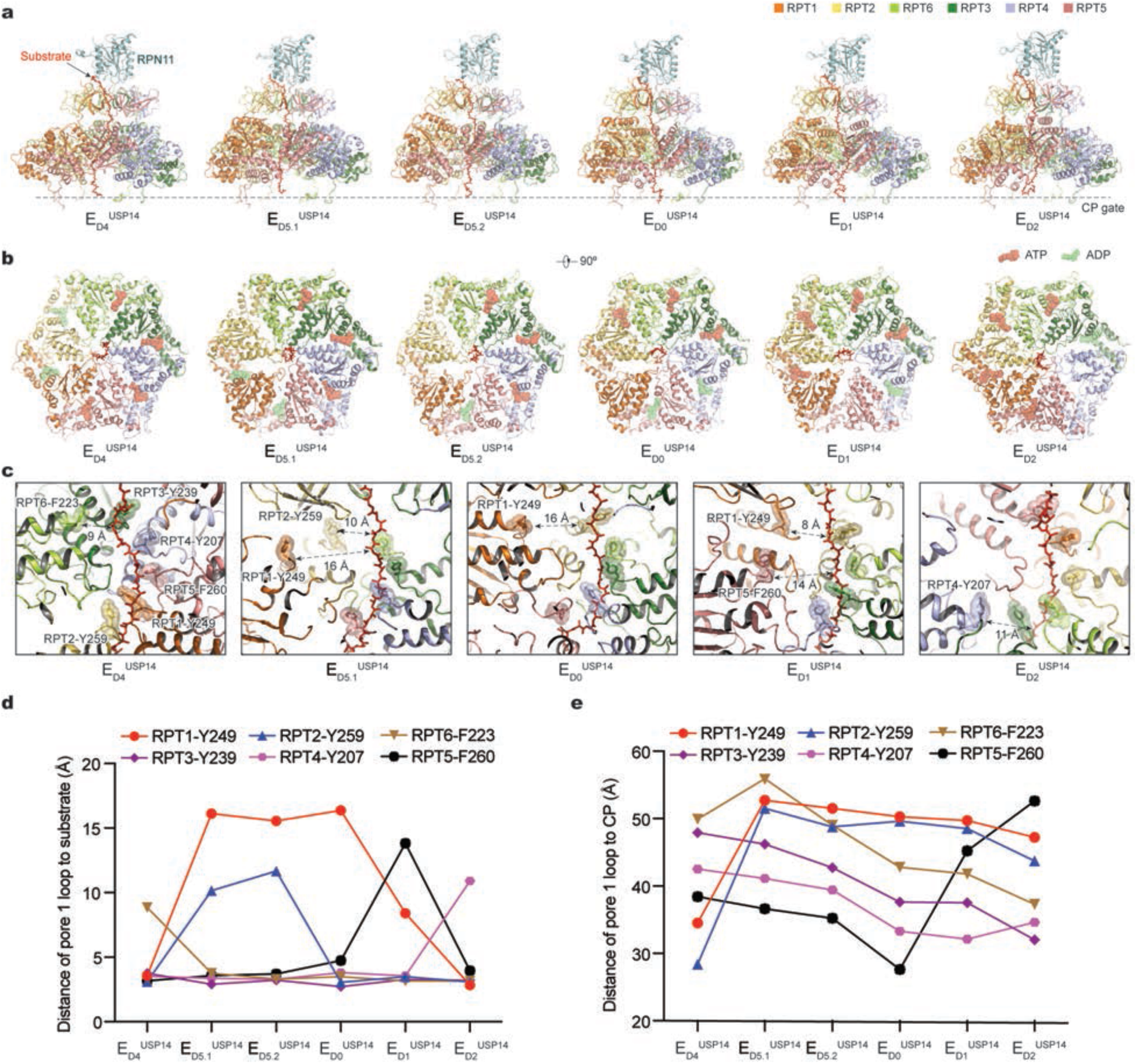
Comparison of the AAA-ATPase motor structures in different USP14-bound states and substrate interactions with the pore-loops. **a**, Side views of the ATPase-RPN11 subcomplex interacting with the substrate in six states from E_D4_^USP14^ to E_D2_^USP14^. The substrate in each state is modelled as a polypeptide backbone structure and is represented with red sticks. RPN11 and the ATPases are rendered as transparent cartoons to show the substrate translocating inside the axial channel. The relative location of the CP is marked by the horizontal dashed line. Top right, color codes of subunits used in all panels. **b**, Corresponding top views of the ATPase motors of six states in (**a**). Nucleotides are shown in stick representation. The sphere representation of ADP and ATP is in green and in red, respectively. The structures are aligned together against their CP components. **c**, Varying architecture of the pore-1 loop staircase interacting with the substrate in five states. State E_D5.2_^USP14^ is not included for its resemblance to state E_D5.1_^USP14^. Aromatic residues in the pore-1 loops are labelled and shown in stick representation superimposed with transparent sphere representation for highlighting. The distances from disengaged pore-1 loops to the substrate are marked. **d**, Plots of distance from the pore-1 loop of each RPT subunit to the substrate in six distinct conformational states. **e**, Plots of distance of the pore-1 loop of each RPT subunit to the CP in six distinct conformational states.

**Extended Data Fig 6.**
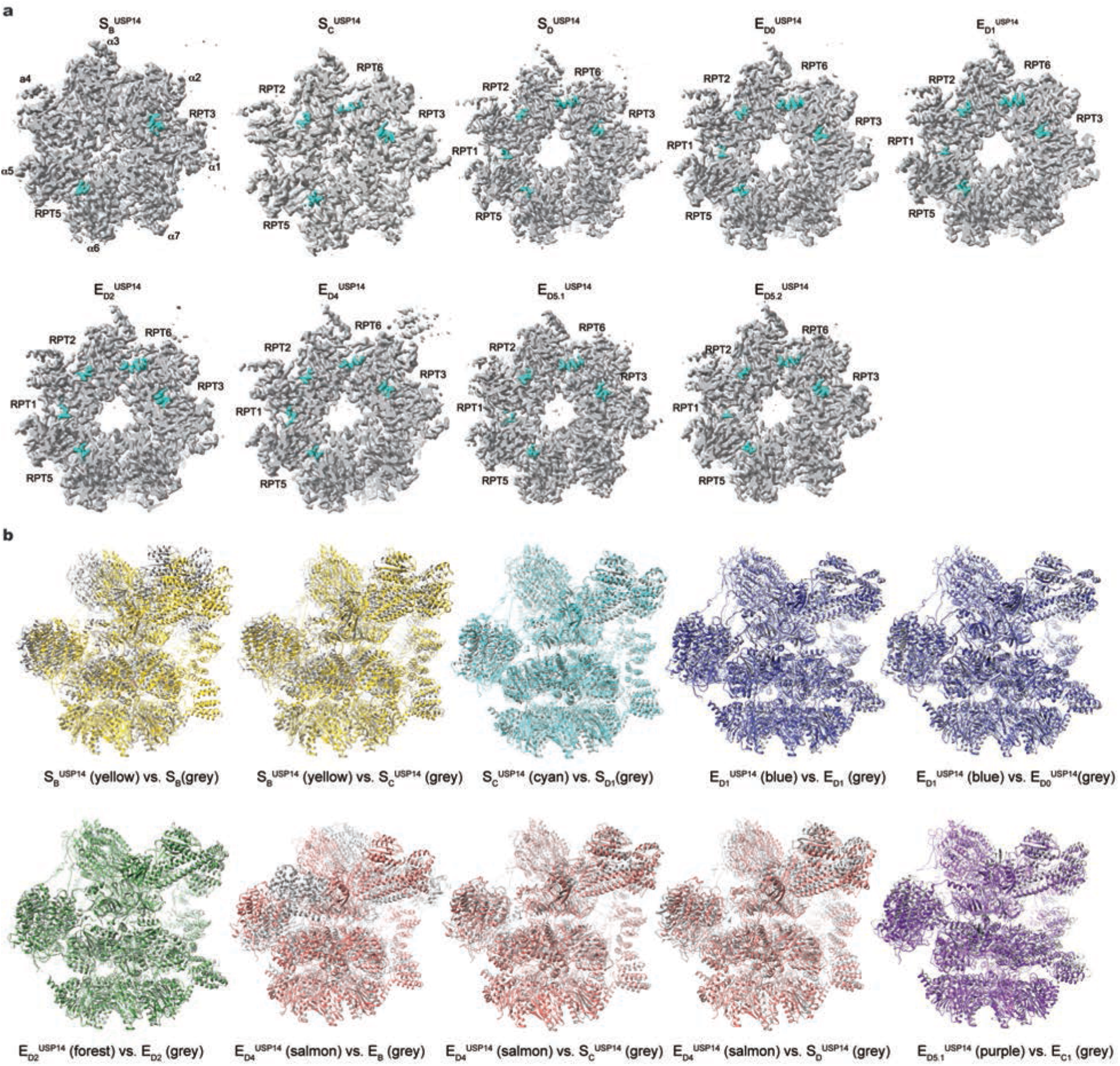
Comparison of the CP and RP between different states. **a**, Comparison of the RP–CP interface and RPT C-terminal tail insertions into the α-pockets of the CP in different states. The cryo-EM densities of the CP subcomplexes are shown as a grey surface representation, while the RPT C-tails are colored green. **b**, Comparison of the RP structures in different states and previously published cryo-EM structures. These structures are aligned together against their CPs. USP14 and ubiquitin are not shown here for clarity in exhibiting differences of the RP structures. Previous structures (PDB ID) used for the comparison include substrate-free states S_B_ (5VFT), S_D1_ (5VFP), and substrate-bound states E_B_ (6MSE), E_C1_ (6MSG), E_D1_ (6MSJ), and E_D2_ (6MSK).

**Extended Data Fig. 7.**
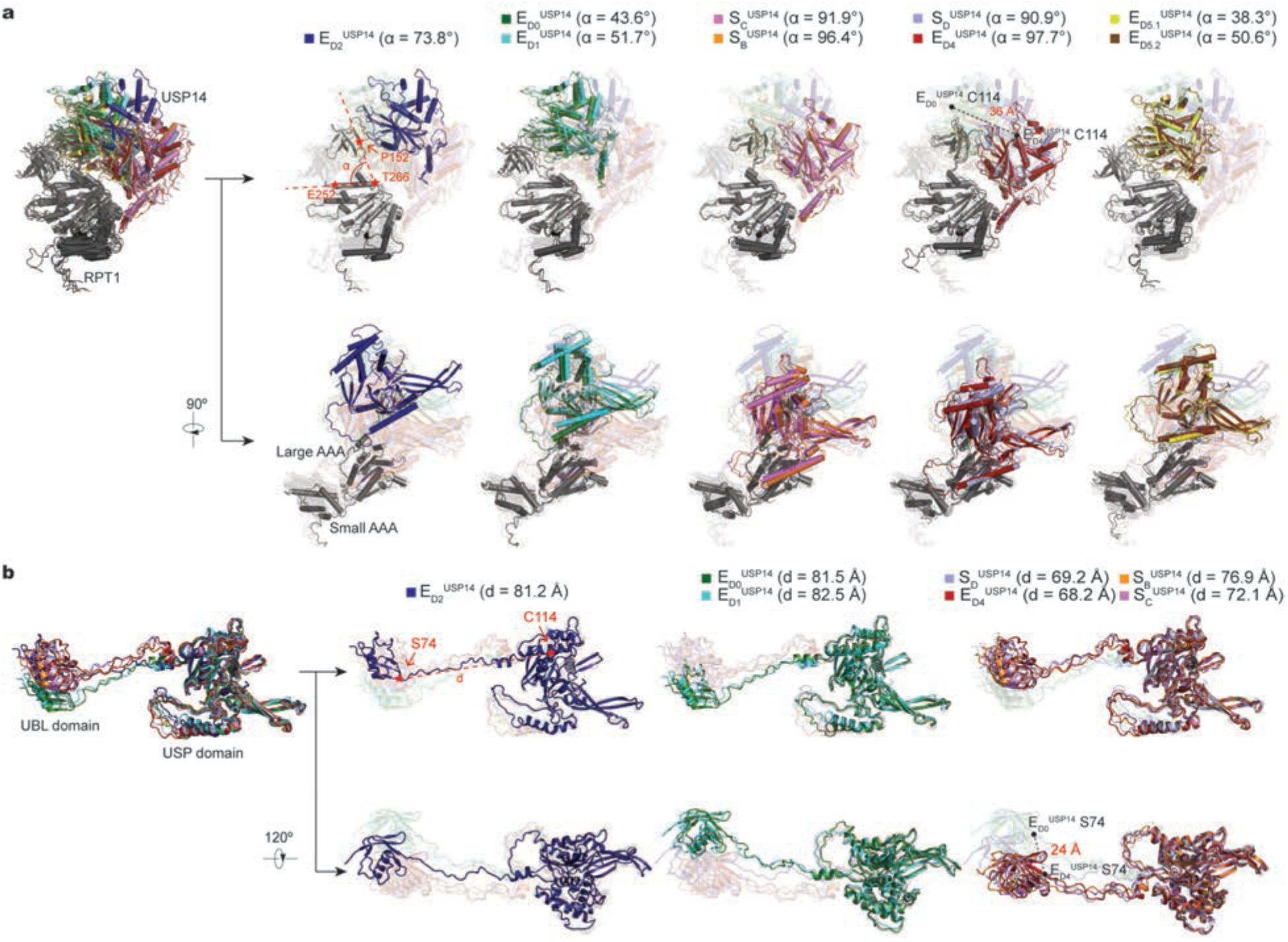
Dynamics of USP14 bound to the proteasome. **a**, Superposition of the USP14-RPT1 subcomplex structures from different states aligned against the RPT1 large AAA subdomain. USP14 rotates together with the RPT1 OB domain and moves up over 36 Å (from E_D4_^USP14^ to E_D0_^USP14^) relative to the RPT1 AAA domain. The angle between the OB domain and the AAA domain is measured and labelled for each state. **b**, Superposition of the USP14 structures from different states aligned against their USP domain. The UBL domain moves up over 24 Å (from E_D4_^USP14^ to E_D0_^USP14^) relative to the USP domain. The distance between S74 and C114 is measured and labelled for each state.

**Extended Data Fig 8.**
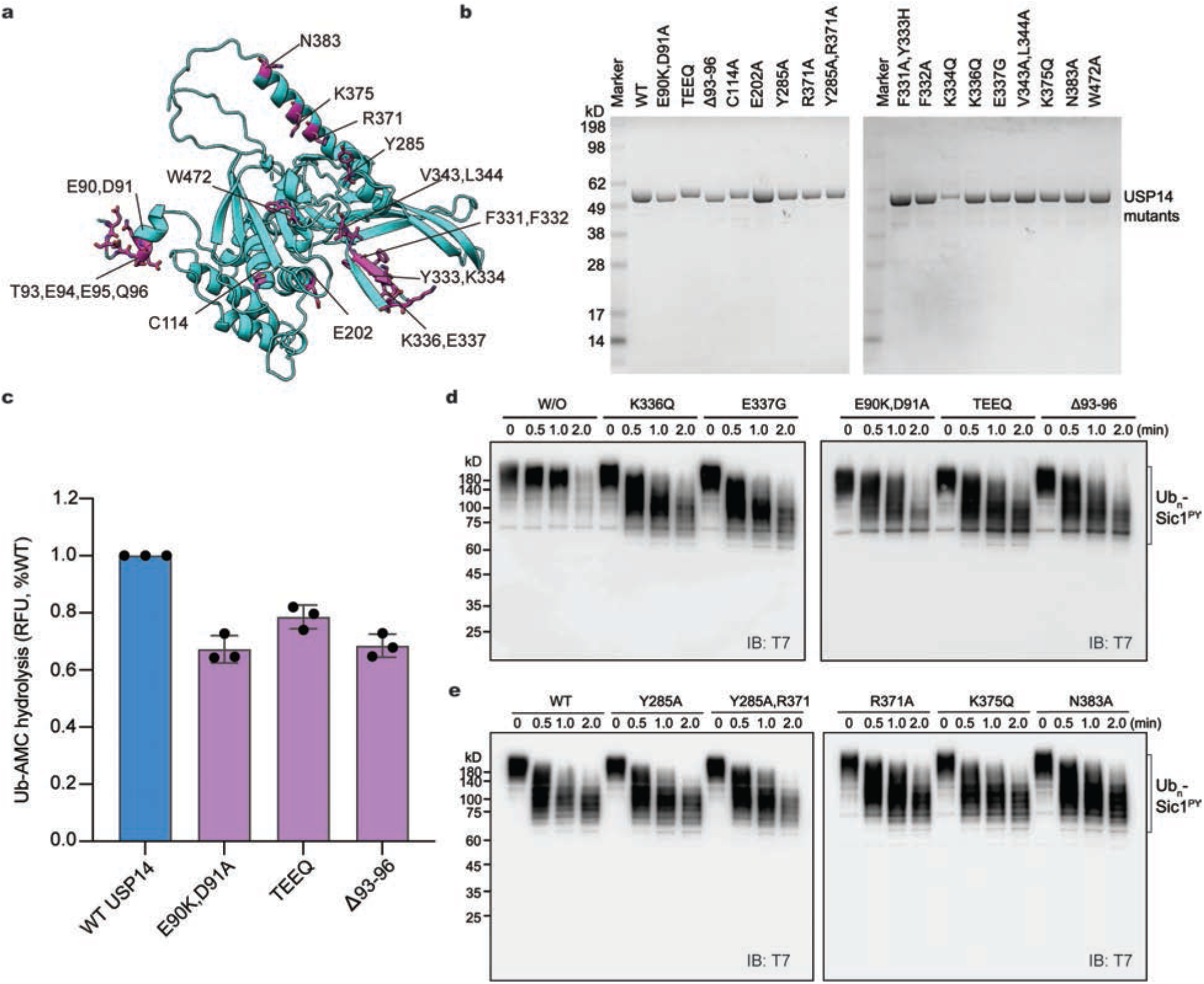
Structure-based site-directed mutagenesis. **a,** Mapping of the potential RPT-binding sites onto the USP14 structure in the E_D4_^USP14^ model. **b,** Purification of USP14 mutants and analyzed by SDS/PAGE and stained with Coomassie blue. **c,** Ub-AMC hydrolysis assay to measure the DUB activity of USP14 mutants in the presence of the human proteasome. Data are presented as mean ± s.d. from 3 independent experiments. **d-e,** *In vitro* degradation of Ub_*n*_-Sic1^PY^ by the 26S proteasome in the presence of USP14 mutants (repeated 3 times). Samples were analyzed by SDS–PAGE/Western blot using anti-T7 antibody. TEEQ, insertion of TEEQ after residue 92. Δ93-96, mutant with deletion of residues 93-96. W/O, the proteasome without binding to USP14. WT, the wildtype USP14-bound proteasome.

**Extended Data Fig. 9.**
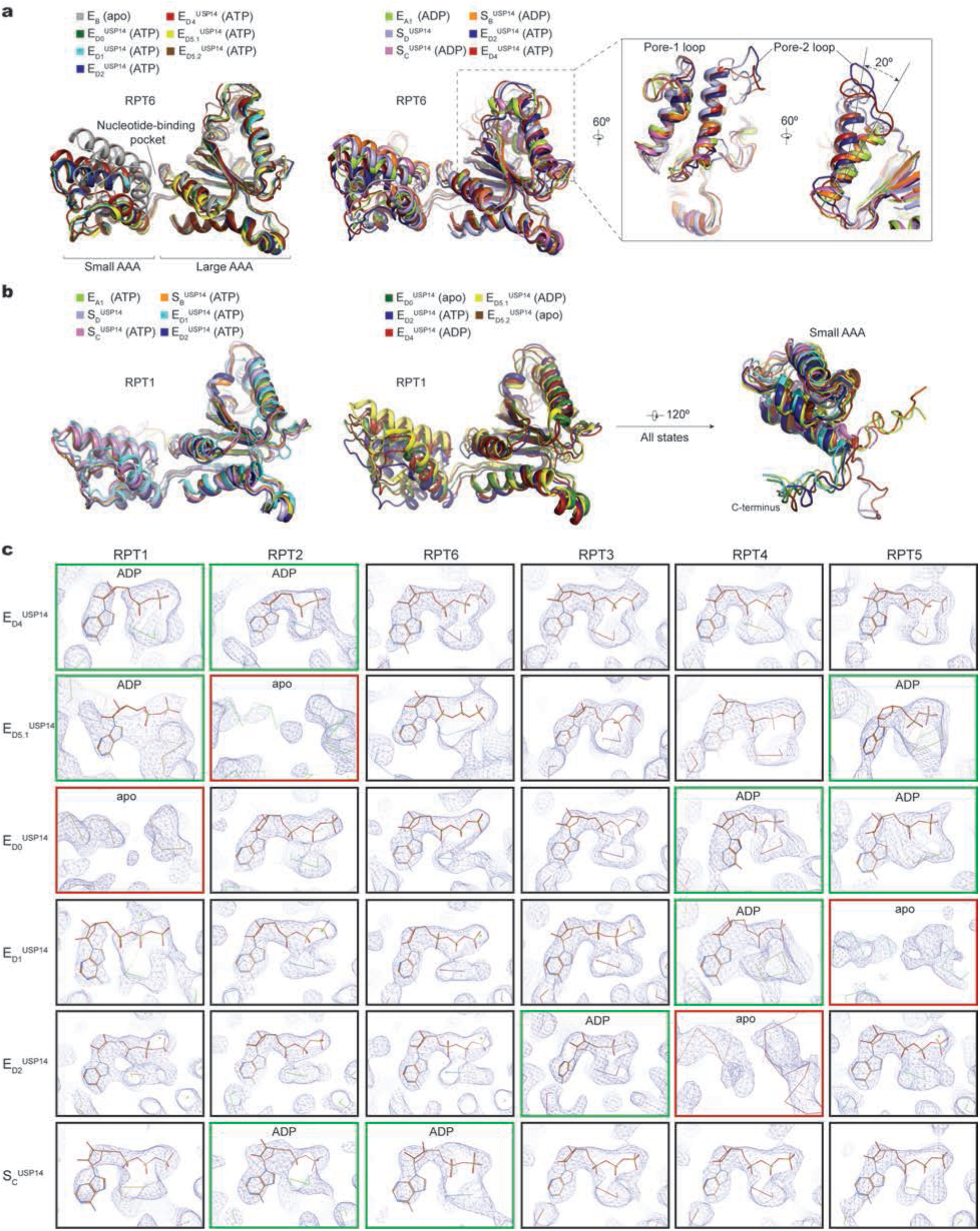
The AAA domain structures and nucleotide states in different USP14-bound proteasome states. **a**, Superposition of the RPT6 AAA domain structures from different states aligned against the large AAA subdomain. Left, comparison of the RPT6 AAA structures in the ATP-bound states and state E_B_ (PDB ID: 6MSE). Middle, comparison of state E_D2_^USP14^, state E_D5.2_^USP14^ and the ADP-bound states shows conformational changes of the AAA domain driven by the ATP hydrolysis. Right, the open-gate states E_D2_^USP14^, E_D4_^USP14^, and S_D_^USP14^ show different refolding of both the pore-1 and pore-2 loops. **b**, Superposition of the RPT1 AAA domain structures from different states aligned against the large AAA subdomain. Left, comparison of the structures in the ATP-bound states. Middle, comparison of the structures in different nucleotide-binding states. Right, the C-tails of RPT6 exhibits three major orientations. **c**, The nucleotide densities fitting with atomic models are shown in blue mesh. All close-up views were directly screen-copied from Coot after atomic modelling into the density maps without modification. At the contour level commonly used for atomic modelling, the potential nucleotide densities in the apo-like subunits mostly disappear, although they can appear as partial nucleotide shapes at a much lower contour level. Other states are not shown here due to the limited local resolution and are hypothetically assigned for nucleotide types or states based on the densities, the openness of corresponding nucleotide-binding pockets as well as their homologous structural models of higher resolution if available.

**Extended Data Table 1.**
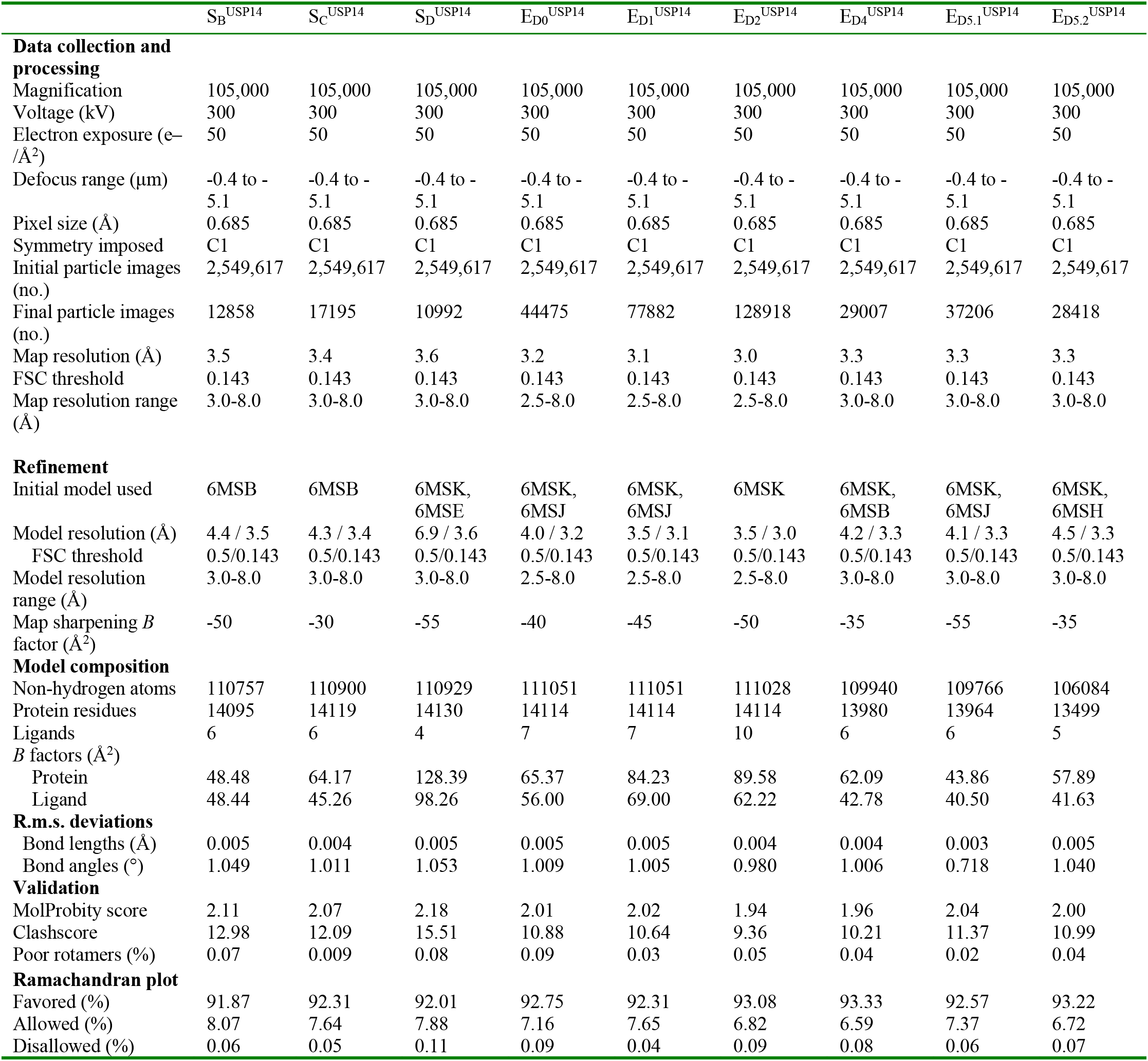
Cryo-EM data collection, refinement and validation statistics.

|  | S<sub>B</sub><sup>USP14</sup> | S<sub>C</sub><sup>USP14</sup> | S<sub>D</sub><sup>USP14</sup> | E<sub>D0</sub><sup>USP14</sup> | E<sub>D1</sub><sup>USP14</sup> | E<sub>D2</sub><sup>USP14</sup> | E<sub>D4</sub><sup>USP14</sup> | E<sub>D5.1</sub><sup>USP14</sup> | E<sub>D5.2</sub><sup>USP14</sup> |
| --- | --- | --- | --- | --- | --- | --- | --- | --- | --- |
| **Data collection and processing** |  |  |  |  |  |  |  |  |  |
| Magnification | 105,000 | 105,000 | 105,000 | 105,000 | 105,000 | 105,000 | 105,000 | 105,000 | 105,000 |
| Voltage (kV) | 300 | 300 | 300 | 300 | 300 | 300 | 300 | 300 | 300 |
| Electron exposure (e<sup>-</sup>/Å<sup>2</sup>) | 50 | 50 | 50 | 50 | 50 | 50 | 50 | 50 | 50 |
| Defocus range (μm) | -0.4 to -5.1 | -0.4 to -5.1 | -0.4 to -5.1 | -0.4 to -5.1 | -0.4 to -5.1 | -0.4 to -5.1 | -0.4 to -5.1 | -0.4 to -5.1 | -0.4 to -5.1 |
| Pixel size (Å) | 0.685 | 0.685 | 0.685 | 0.685 | 0.685 | 0.685 | 0.685 | 0.685 | 0.685 |
| Symmetry imposed | C1 | C1 | C1 | C1 | C1 | C1 | C1 | C1 | C1 |
| Initial particle images (no.) | 2,549,617 | 2,549,617 | 2,549,617 | 2,549,617 | 2,549,617 | 2,549,617 | 2,549,617 | 2,549,617 | 2,549,617 |
| Final particle images (no.) | 12858 | 17195 | 10992 | 44475 | 77882 | 128918 | 29007 | 37206 | 28418 |
| Map resolution (Å) | 3.5 | 3.4 | 3.6 | 3.2 | 3.1 | 3.0 | 3.3 | 3.3 | 3.3 |
| FSC threshold | 0.143 | 0.143 | 0.143 | 0.143 | 0.143 | 0.143 | 0.143 | 0.143 | 0.143 |
| Map resolution range (Å) | 3.0-8.0 | 3.0-8.0 | 3.0-8.0 | 2.5-8.0 | 2.5-8.0 | 2.5-8.0 | 3.0-8.0 | 3.0-8.0 | 3.0-8.0 |
| **Refinement** |  |  |  |  |  |  |  |  |  |
| Initial model used | 6MSB | 6MSB | 6MSK, 6MSE | 6MSK, 6MSJ | 6MSK, 6MSJ | 6MSK | 6MSK, 6MSB | 6MSK, 6MSJ | 6MSK, 6MSH |
| Model resolution (Å) | 4.4 / 3.5 | 4.3 / 3.4 | 6.9 / 3.6 | 4.0 / 3.2 | 3.5 / 3.1 | 3.5 / 3.0 | 4.2 / 3.3 | 4.1 / 3.3 | 4.5 / 3.3 |
| FSC threshold | 0.5/0.143 | 0.5/0.143 | 0.5/0.143 | 0.5/0.143 | 0.5/0.143 | 0.5/0.143 | 0.5/0.143 | 0.5/0.143 | 0.5/0.143 |
| Model resolution range (Å) | 3.0-8.0 | 3.0-8.0 | 3.0-8.0 | 2.5-8.0 | 2.5-8.0 | 2.5-8.0 | 3.0-8.0 | 3.0-8.0 | 3.0-8.0 |
| Map sharpening *B* factor (Å<sup>2</sup>) | -50 | -30 | -55 | -40 | -45 | -50 | -35 | -55 | -35 |
| **Model composition** |  |  |  |  |  |  |  |  |  |
| Non-hydrogen atoms | 110757 | 110900 | 110929 | 111051 | 111051 | 111028 | 109940 | 109766 | 106084 |
| Protein residues | 14095 | 14119 | 14130 | 14114 | 14114 | 14114 | 13980 | 13964 | 13499 |
| Ligands | 6 | 6 | 4 | 7 | 7 | 10 | 6 | 6 | 5 |
| *B* factors (Å<sup>2</sup>) |  |  |  |  |  |  |  |  |  |
| Protein | 48.48 | 64.17 | 128.39 | 65.37 | 84.23 | 89.58 | 62.09 | 43.86 | 57.89 |
| Ligand | 48.44 | 45.26 | 98.26 | 56.00 | 69.00 | 62.22 | 42.78 | 40.50 | 41.63 |
| **R.m.s. deviations** |  |  |  |  |  |  |  |  |  |
| Bond lengths (Å) | 0.005 | 0.004 | 0.005 | 0.005 | 0.005 | 0.004 | 0.004 | 0.003 | 0.005 |
| Bond angles (°) | 1.049 | 1.011 | 1.053 | 1.009 | 1.005 | 0.980 | 1.006 | 0.718 | 1.040 |
| **Validation** |  |  |  |  |  |  |  |  |  |
| MolProbity score | 2.11 | 2.07 | 2.18 | 2.01 | 2.02 | 1.94 | 1.96 | 2.04 | 2.00 |
| Clashscore | 12.98 | 12.09 | 15.51 | 10.88 | 10.64 | 9.36 | 10.21 | 11.37 | 10.99 |
| Poor rotamers (%) | 0.07 | 0.009 | 0.08 | 0.09 | 0.03 | 0.05 | 0.04 | 0.02 | 0.04 |
| **Ramachandran plot** |  |  |  |  |  |  |  |  |  |
| Favored (%) | 91.87 | 92.31 | 92.01 | 92.75 | 92.31 | 93.08 | 93.33 | 92.57 | 93.22 |
| Allowed (%) | 8.07 | 7.64 | 7.88 | 7.16 | 7.65 | 6.82 | 6.59 | 7.37 | 6.72 |
| Disallowed (%) | 0.06 | 0.05 | 0.11 | 0.09 | 0.04 | 0.09 | 0.08 | 0.06 | 0.07 |

**Extended Data Table 2.**
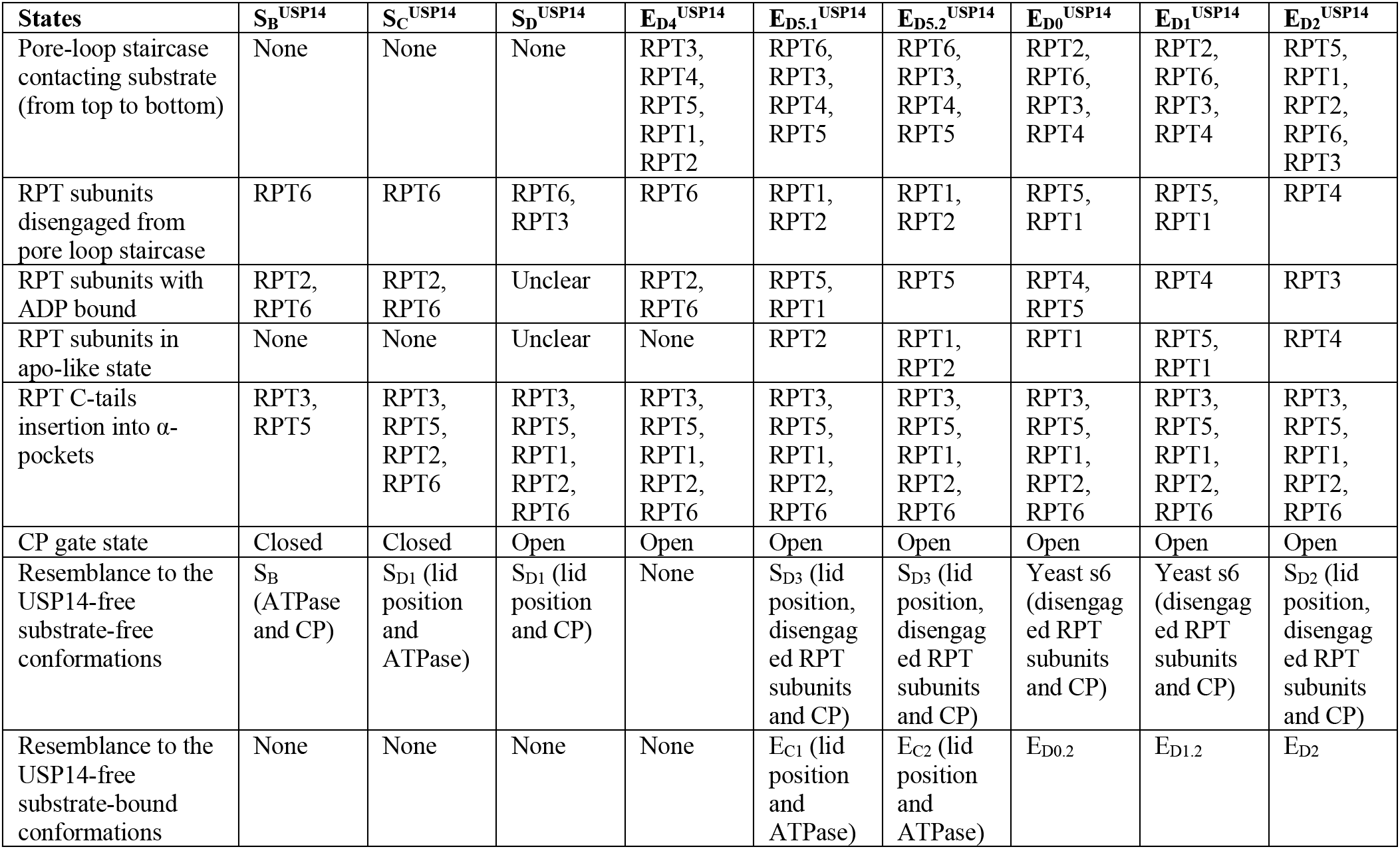
Summary of key structural features of USP14-bound states. The nucleotide states in the substrate-engaged state share common features consistent with the previously reported results^10^. ATP is generally bound to the substrate-engaged ATPases above the penultimate contact along the substate. ADP is bound to the ATPase subunit at the lowest position in contact with substrate, which is poised for dissociation from substrate, whereas an apo-like or ADP-bound state is generally found in the substrate-disengaged ATPase subunits that is in the seam of the lock-washer-like ATPase ring.

